# Recognizing dUTPase as a mitotic factor essential for early embryonic development

**DOI:** 10.64898/2026.01.16.699898

**Authors:** Nikolett Nagy, Otília Tóth, Eszter Oláh, László Henn, Gergely Attila Rácz, Edit Szabó, György Várady, Fanni Beatrix Vigh, Zita Réka Golács, Martin Urbán, Tímea Pintér, Orsolya Ivett Hoffmann, László Hiripi, Hilde Loge Nilsen, Angéla Békési, Miklós Erdélyi, Elen Gócza, Gergely Róna, Judit Tóth, Beáta G. Vértessy

## Abstract

dUTPase is universally regarded as a metabolic sanitizing enzyme that protects genomes by preventing the incorporation of uracil into DNA. Despite its essentiality across eukaryotes, no function beyond nucleotide sanitization has been demonstrated. Here, we uncover a conserved, non-canonical role for dUTPase as a regulator of mitosis. Using *Drosophila* and mouse models, we demonstrate that dUTPase loss causes early embryonic lethality characterized by severe mitotic failure that, cannot be rescued by disabling uracil-DNA repair, uncoupling dUTPase essentiality from DNA repair pathways. Mechanistically, dUTPase dynamically associates with the mitotic spindle and centrosomes, and its depletion induces centrosome amplification and chromosome segregation defects. Beyond cell division, dUTPase dosage bidirectionally controls cell migration, linking its mitotic function to cellular behaviors relevant for metastasis. Together, our findings redefine dUTPase as a moonlighting mitotic factor that coordinates centrosome integrity and spindle dynamics, expanding its known repertoire beyond nucleotide metabolism.

**Highlights:**

dUTPase deficiency leads to mitotic defects during early embryonic development in both *Drosophila* and mouse models
dUTPase shows dynamic spatiotemporal localization associated with microtubules through mitosis
dUTPase is essential for the normal number and intracellular localization of centrosomes
dUTPase deficiency counteracts while its overexpression enhances cell migration

## Introduction

The enzyme deoxyuridine-5’-triphosphate nucleotidohydrolase (dUTPase, DUT) is traditionally recognized as a key guardian of genomic integrity. It achieves this by hydrolyzing dUTP to dUMP, thus maintaining a low intracellular dUTP/dTTP ratio [1]. This action minimizes the risk of uracil incorporation into DNA, which can occur due to the inability of most DNA polymerases to discriminate between dUTP and dTTP. In addition to misincorporation, uracil can appear in DNA through spontaneous cytosine deamination, the most common and physiologically relevant DNA lesion. Under normal conditions, this reaction introduces approximately 100-200 uracil bases per diploid human cell per day [2]. The real biological load of uracil *in vivo* can be much higher [3]. These uracil residues, if not corrected, are a major source of C-to-T transition mutations, contributing to genome instability, cancer development, and long-term evolutionary change [4].

To counteract these threats, cells rely on uracil-DNA glycosylases (UDGs), such as UNG and SMUG1, to excise uracil residues, initiating base excision repair (BER) [3]. The enzyme dUTPase has dual importance in uracil exclusion and nucleotide biosynthesis, as it decreases dUTP levels and provides dUMP as a precursor for *de novo* thymidylate (dTMP) biosynthesis [1]. This function is particularly important during cell proliferation, where demand for nucleotide precursors is high; thus, mRNA and protein expression of dUTPase was found to be elevated in proliferative tissues [5]–[8] and in cancer cells [9], [10]. The enzyme is encoded by the *DUT* gene in mammals, and through alternative splicing and promoter usage, nuclear, mitochondrial, and cytoplasmic variants can be generated [8], [11], [12] . Moreover, phosphorylation of nuclear dUTPase during the G2/M phase of the cell cycle [13], [14], [15] indicates regulated activity during mitosis.

Although the metabolic and dNTP-sanitizing role of dUTPase has been extensively characterized over the past four decades, this canonical function may obscure additional, potentially more consequential cellular roles. Importantly, in every multicellular organism examined to date that encodes dUTPase, the protein has proven indispensable for viability [16]–[19]. Genetic depletion or knockout of *DUT* results in lethality in diverse systems, including *Arabidopsis thaliana* [20], [21], *Trypanosoma brucei* [22], *Saccharomyces cerevisiae* [16], [23], *Caenorhabditis elegans* [24], *Drosophila melanogaster* [25], and human cells [26], [27]. Lethality cannot be rescued even by manipulating downstream uracil repair pathways [24], [28]–[33].

Importantly, the developmental timing of lethality varies among organisms, pointing to context-dependent requirements for dUTPase. In *Drosophila*, dUTPase knockdown triggers developmental arrest at the pupal stage, even though early larval stages can transiently tolerate its absence [25]; however, during early embryonic development, maternal dUTPase is provided [34]. To determine whether dUTPase is required for early embryonic development, the maternal pool of dUTPase would need to be eliminated, an issue that has not yet been addressed in the literature. In mice, complete *Dut* knockout results in early embryonic lethality with perturbed blastocyst growth [18], suggesting a non-redundant role for dUTPase during the earliest stages of mammalian development. Recently, it was demonstrated in zebrafish that disruption of the *dut* gene leads to strong lethality during post-embryonic development, where maternal supply is exhausted [19]. In contrast, elimination of other dNTP pool-sanitizing enzymes, most notably members of the NUDIX hydrolase family and SAMHD1, does not result in lethality, and these enzymes are not strictly required for normal development [35], [36], [37]. Several additional observations further challenge the view that dUTPase functions solely as a metabolic sanitizing enzyme, e.g., the existence of multiple dUTPase isoforms with distinct subcellular localization and regulatory phosphorylation during the cell cycle. Additional findings, including roles in nuclear trafficking regulation [14], [15], and phenotypes of catalytically inactive mutants [16], [23], [28]–[30], further suggest that dUTPase may participate in processes tied to the cell cycle, DNA damage response, and chromatin organization. Together, these observations argue for a more nuanced, context-dependent essentiality of dUTPase in metazoan biology.

This study aims to re-evaluate the cellular functions of dUTPase using multiple model organisms and state-of-the-art experimental approaches, with a specific focus on its underexplored roles in the mammalian cell cycle. By shifting the perspective beyond uracil metabolism, we sought to uncover novel aspects of dUTPase biology that may be essential for cell division, genome stability, or developmental regulation. Here, we examined the consequences of dUTPase loss during early embryonic development in *Drosophila melanogaster* using CRISPR gene editing as well as RNAi-mediated silencing to eliminate both zygotic and maternal sources. We demonstrated that elimination of maternal dUTPase caused mitotic defects that led to early embryonic lethality. Furthermore, in mice, *Dut* knockout could not be rescued by depletion of enzymes involved in uracil repair, Ung and Smug1, resulting in early embryonic lethality. Immunofluorescence analysis revealed that dUTPase exhibits spatiotemporal and cell type-specific localization during embryonic development. In human cells, dUTPase dynamically associates with the mitotic spindle and centrosomes, and its shRNA-mediated depletion induces centrosome amplification and mitotic defects. Notably, dUTPase deficiency counteracts, while its overexpression enhances the cell migration rate. Together, these findings highlight previously unrecognized roles of dUTPase that may also have implications for its therapeutic targeting, particularly in the treatment of cancer.

## Results and Discussion

### Loss of dUTPase in *Drosophila melanogaster* leads to mitotic defects and lethality

In *Drosophila melanogaster*, a key model organism in developmental biology, the lethal effect of dUTPase deficiency has been previously reported only during metamorphosis [25]. In this study, we sought to characterize dUTPase depletion during *Drosophila* development using CRISPR gene editing and shRNA-mediated silencing. Phenotype analysis of the mutant line generated by CRISPR gene editing resulted in an 867 bp deletion, while silencing of dUTPase was achieved by maternal depletion of gene product (RNAi) targeting the 3’-UTR regions (Figure 1A). In *Drosophila*, the *dut* gene encodes two isoforms of dUTPase, a nuclear, 23 kDa (Dut-N) isoform containing an NLS sequence and a cytoplasmic, 21 kDa (Dut-C) isoform [34], which were used as rescue constructs (Figure 1A).

**Figure 1.**
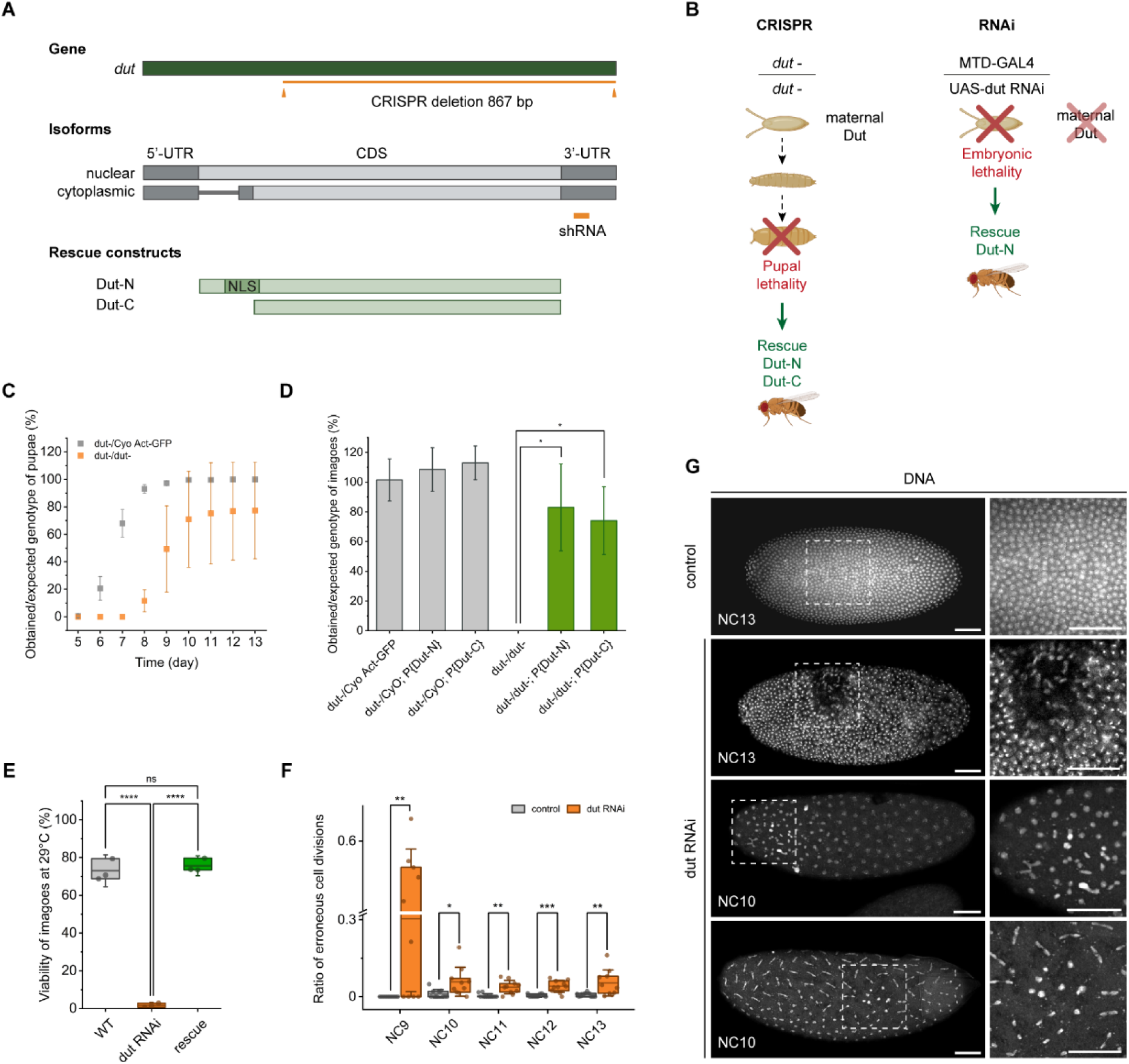
Loss of dUTPase in *Drosophila melanogaster* leads to mitotic defects and lethality. (A) Schematic representation of the *dut* gene (dark green) and isoforms (gray) in *Drosophila melanogaster.* Target sites of CRISPR gene editing and shRNA silencing are shown in orange. 5’- and 3’-UTR regions are shown in dark gray. Rescue constructs of nuclear (Dut-N) and cytoplasmic (Dut-C) isoforms of dUTPase are shown in light green. NLS: nuclear localization signal. CDS: coding sequence. (B) Schematic representation of dUTPase manipulation using CRISPR or RNAi. Knockout of *dut* by CRISPR gene editing leads to pupal lethality and can be rescued by either Dut-N or Dut-C isoforms with their shared native promoter. Maternally driven silencing (MTD-GAL4/UAS-*dut* RNAi) of *dut* leads to embryonic lethality, which can be rescued with Dut-N transgene with its native promoter. (C) Pupariation dynamics of heterozygous (*dut-*/Cyo Act-GFP, gray) and homozygous knockout (*dut-/dut-*, orange) larvae generated by CRISPR gene editing. Obtained/expected genotype ratios were determined based on the number of counted heterozygote animals. Whiskers show standard deviation. (D) Percentage of obtained/expected hatched adults of heterozygous (gray), knockout, and rescue (green) genotypes as compared to heterozygous animals generated by CRISPR gene editing. Whiskers show standard deviation. Statistical analysis was carried out using Welch’s t-test (*: p<0.04). (E) Viability of dUTPase knockdown (*dut* RNAi) animals (orange) at 29°C compared to WT (gray) and Dut-N rescue (green) imagoes. Whiskers show standard deviation. One-way ANOVA with Tukey multiple comparisons test was used (****: p<0.0001, ns: non-significant). (F) Ratio of erroneous cell divisions from NC9 to NC13 in WT (gray) and *dut* RNAi (orange) embryos. NC: nuclear cycle. Axis break is set from 0.32 to 0.5. Whiskers show standard deviation. Statistical analysis was carried out using Kruskal-Wallis test with Dunn’s multiple comparisons test (*: p<0.015, **: p<0.006, ***: p<0.001). (G) Microscopy images of NC13 control and NC13 and NC10 *dut* RNAi *Drosophila* embryos. Framed sections showing mitotic defects are enlarged. Scale bar 50 μm.

CRISPR knockout of dUTPase resulted in pupal lethality, which can be rescued by either Dut-N or Dut-C isoforms (Figure 1B, Supplementary Figure S1A). Although ∼80% of the knockout animals reach the pupal stage, they exhibit altered dynamics, a decreased pupariation rate, and are unable to develop further than the pupal stage, as compared to heterozygous flies (Figure 1C). Either Dut-N or Dut-C isoforms expressed under their native promoter significantly increased the number of obtained fully developed adults compared to dUTPase knockout animals, where no flies could hatch (Figure 1D). To verify the nuclear and cytoplasmic localization of Dut-N and Dut-C rescue constructs, respectively, we applied immunofluorescence in the ovaries of mutant animals (Supplementary Figure S1B).

Since maternal nurse cells supply the early embryo with abundant proteins and mRNAs, including dUTPase [34], we aimed to investigate the effect of dUTPase loss during early embryonic development. Therefore, we used maternally driven RNA interference (MTD-GAL4/UAS-*dut* RNAi) to deplete maternal dUTPase contribution and establish a dUTPase-deficient embryonic environment (Supplementary Figure S1C). We found that maternal depletion caused severe embryonic lethality; however, this effect was fully restored by transgenic Dut-N expressed under its native promoter (Figure 1B and E). Lethality was temperature-dependent: the MTD-GAL4/UAS-*dut* RNAi construct was more effective at 29°C, causing near-100% embryonic lethality (Figure 1E), whereas at 25°C, lethality was lower and distributed across developmental stages: ∼50% of embryos, ∼10% of larvae, and ∼15% of pupae failed to survive (Supplementary Figure S1D and E). It is well known that GAL4-driven silencing is more efficient at elevated temperatures – our results, showing higher lethality at 29°C than at 25°C, agree with this. Video microscopy of control (Supplementary Video S1) and dUTPase-depleted (Supplementary Video S2) embryos revealed disrupted early syncytial divisions. Analyzing the frequency of erroneous cell divisions during embryonic development, we observed a significantly higher ratio of mitotic defects in dUTPase RNAi than in control embryos across NC9 to NC13, particularly during the early NC9 phase (Figure 1F). Microscopy images show disrupted early syncytial divisions, frequent chromosome bridges, and abnormal chromosomal structures observed in NC10 and NC13 dUTPase RNAi embryos (Figure 1G).

RNAi silencing of dUTPase results in embryonic lethality when no maternal dUTPase is present, and mRNA levels are highly elevated at the early embryonic stage (Supplementary Figure S1F), suggesting that dUTPase is indispensable for *Drosophila* embryogenesis, with its absence leading to mitotic failure. As demonstrated previously, the mRNA level of dUTPase decreases during the larval stage [25], enabling knockout animals to survive. Interestingly, in most larval cells, such as in the salivary gland, fat body, and digestive system, endoreplication occurs, during which DNA repeatedly duplicates without cell division in the absence of a canonical mitotic apparatus, resulting in polyploidy [38], where dUTPase mRNA level is low according to modENCODE mRNA data available on FlyBase. Although DNA synthesis occurs continuously during endoreplication, and according to the literature, dUTPase is important for maintaining a low dUTP/dTTP ratio during S phase, these cells lack dUTPase at both the mRNA and protein levels. In contrast, dUTPase is highly expressed in larval imaginal discs and central nervous system (CNS) [5], where normal mitosis occurs. Interestingly, it has been previously demonstrated in S2 *Drosophila* cells that dUTPase isoforms exhibit dynamic localization during cell division [39]. The eYFP-tagged isoforms of dUTPase were microinjected into *Drosophila* embryos, and their localization was followed with fluorescence microscopy. Reevaluation of these results, focusing on mitotic cells, suggests that both isoforms of dUTPase localize to the microtubule network during mitosis.

In conclusion, these data suggest that dUTPase might have a role in mitotic processes. To assess whether this phenomenon is conserved and to extend the relevance of our observations, we investigated dUTPase function in a mammalian model as well.

### Knockout of *Dut* in mice cannot be rescued by *Ung* or *Smug1* deletion

A previous study has shown that *Dut^-/-^* mouse embryos die shortly after implantation [18]. In contrast, single knockouts of *Ung* or *Smug1*, as well as double knockout *Ung^-/-^ Smug1^-/-^* mice, are viable and fertile, despite having elevated genomic uracil levels [3]. We were interested in whether the *Dut^-/-^* embryos can be rescued by the depletion of base excision repair (BER) enzymes, Ung and Smug1. Therefore, we crossed *Ung^-/-^*, *Smug1^-/-^,* or *Ung^-/-^ Smug1^-/-^* mice with *Dut^+/-^*animals.

Neither *Ung^-/-^ Dut^-/-^* nor *Smug1^-/-^ Dut^-/-^* (Supplementary Figure S2A and B) or *Ung^-/-^ Smug1^-/-^ Dut^-/-^* (Figure 2A and B) embryos were viable. Furthermore, microscopy analysis of 3.5 dpc (days post coitum) embryos revealed that *Ung^-/-^ Smug1^-/-^ Dut^-/-^* embryos contained fewer cells than their *Dut^+/-^* and *Dut^+/+^* counterparts (Figure 2C, Supplementary Figure S2A). The unexpected lethality of the triple knockout animals led us to hypothesize about a hitherto unknown function of dUTPase during early development. To elucidate this potential function, we analyzed the overall localization pattern of dUTPase in early embryonic stages of wild-type (WT) animals.

**Figure 2.**
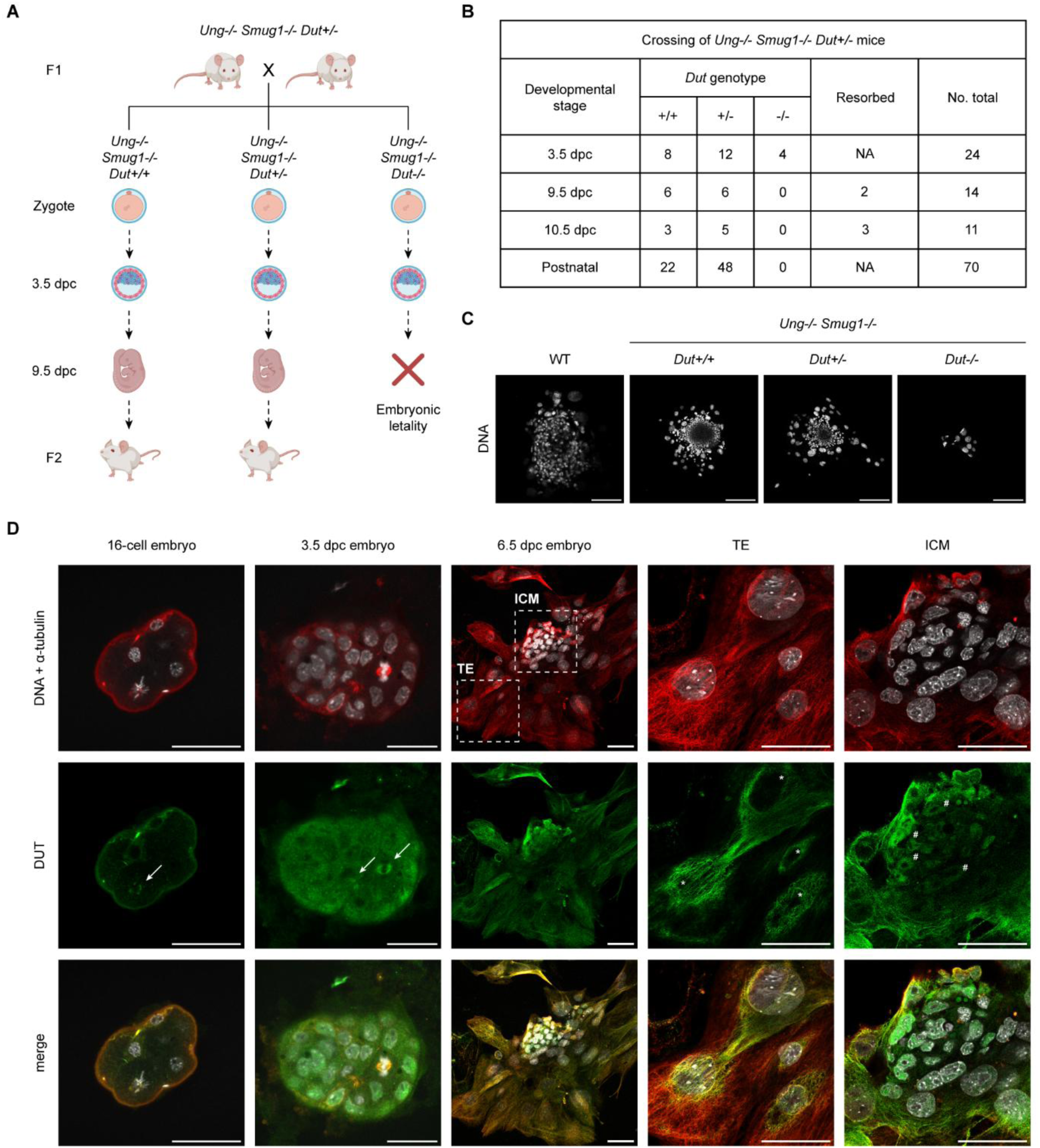
Investigation of *Dut* knockout and WT mouse embryos. (A) Schematic representation of crossings of *Ung^-/-^ Smug1^-/-^ Dut*^+/-^ genotype mice. dpc: day post coitum. (B) Genotyping results of offspring from crossings of *Ung^-/-^ Smug1^-/-^ Dut^+/-^* mice. (C) Microscopy analysis of WT embryo and embryos from *Ung^-/-^ Smug1^-/-^ Dut^+/-^* crossings at 3.5 dpc. DNA is shown in gray. Scale bar 100 μm. (D) Immunostaining of DNA (gray), α-tubulin (red), and dUTPase (green) in WT 16-cell, 3.5 dpc, and 6.5 dpc embryos. Framed inlets showing trophectoderm (TE) and inner cell mass (ICM) are magnified in separate panels. Scale bar 50 μm. Arrows indicate mitotic and central spindles in dividing cells. Asterisks highlight the absence of dUTPase in nuclei of TE, while hashtags indicate nuclear presence of dUTPase in ICM.

### dUTPase is localized to the spindle apparatus in the mouse embryo

To investigate the role of dUTPase during mouse embryonic development, we identified the localization in 16-cell, early (3.5 dpc), and late (6.5 dpc) blastocyst embryos using immunocytochemistry (Figure 2D, Supplementary Figure S2C). Secondary antibody controls were used to verify the specificity of the immunostaining (Supplementary Figure S2D). Interestingly, the localization of dUTPase exhibited dynamic changes at different embryonic stages. In the early 16-cell stage, dUTPase was not enriched in the nucleus, but a homogenous cellular distribution was observed; however, during cell division, it accumulates on the acentrosomal microtubule-organizing centers (MTOCs) (Figure 2D). In rodents, during embryonic development, transition from acentrosomal to centrosomal spindle formation occurs during pre-implantation development before the blastocyst stage [40], [41]. Notably, according to our previous results, *Dut^-/-^*genotype embryos die shortly after implantation. In 3.5 dpc embryos, dUTPase exhibits homogenous distribution, and in mitotic cells, colocalization with α-tubulin in the mitotic and central spindle can be observed (Figure 2D, Supplementary Figure S2C). In 6.5 dpc blastocysts, dUTPase localization differs across cell types, with nuclear localization in the inner cell mass (ICM) and filamentous pattern in the trophectoderm (TE) cells, associated with α-tubulin. Interestingly, in the TE cells, no nuclear accumulation of dUTPase can be observed (Figure 2D, Supplementary Figure S2C). Previously, detailed analysis of pre-implantation *Dut^-/-^* embryos revealed perturbed growth of both inner cell mass (ICM) and trophectoderm (TE) [18], highlighting the importance of dUTPase in different cell types during embryonic development. In conclusion, we observed a dynamic distribution of dUTPase during early embryonic development, with accumulation in regions where microtubules are present.

### Localization of dUTPase in the different stages of mitosis in human cells

To broaden the scope of our results to human cells, we examined the subcellular localization of dUTPase (Figure 3A) throughout the different phases of mitosis using immunostaining in the HCT116 human colorectal cancer cell line. Consistent findings were obtained in HEK293, U2OS, and MEF cells (Supplementary Figure S3A), confirming that the localization pattern is observed across mammalian cell types. During interphase, the microtubule network stained for α-tubulin exhibited its characteristic filamentous cytoplasmic distribution, while dUTPase displayed predominant nuclear localization with additional cytoplasmic presence and accumulation in distinct foci. As cells entered prophase, dUTPase was excluded from the nucleus and displayed a more homogeneous distribution, consistent with previous reports assessing the phosphorylation of dUTPase [14], which might be linked to mitotic entry. From prometaphase through metaphase, dUTPase is prominently localized to the forming mitotic spindle, colocalizing with α-tubulin (Figure 3A, Supplementary Figure S3A). This dynamic redistribution toward the spindle apparatus implies a potential role in spindle assembly or stability. In anaphase, enrichment of dUTPase can be observed along both mitotic and central spindle microtubules. In telophase, it localizes to the central spindle and accumulates near the centrosomes. During cytokinesis, dUTPase was observed in the reforming nuclei and centrosomes. Collectively, colocalization of dUTPase with α-tubulin can be observed in the mitotic spindle during prometaphase, metaphase, and anaphase, and in the central spindle during anaphase and telophase (Figure 3A, Supplementary Figure S3A). This transition suggests that dynamic relocalization of dUTPase may accompany mitotic exit and reestablishment of nuclear function. To validate these observations, we generated a stable HCT116 cell line harboring a doxycycline-inducible transgene containing a fluorescent dUTPase fusion protein (DUT-N-eGFP). Visualization of DUT-N-eGFP in mitotic cells (Figure 3B) confirmed its accumulation in the mitotic spindle, underscoring the robustness of this localization pattern.

**Figure 3.**
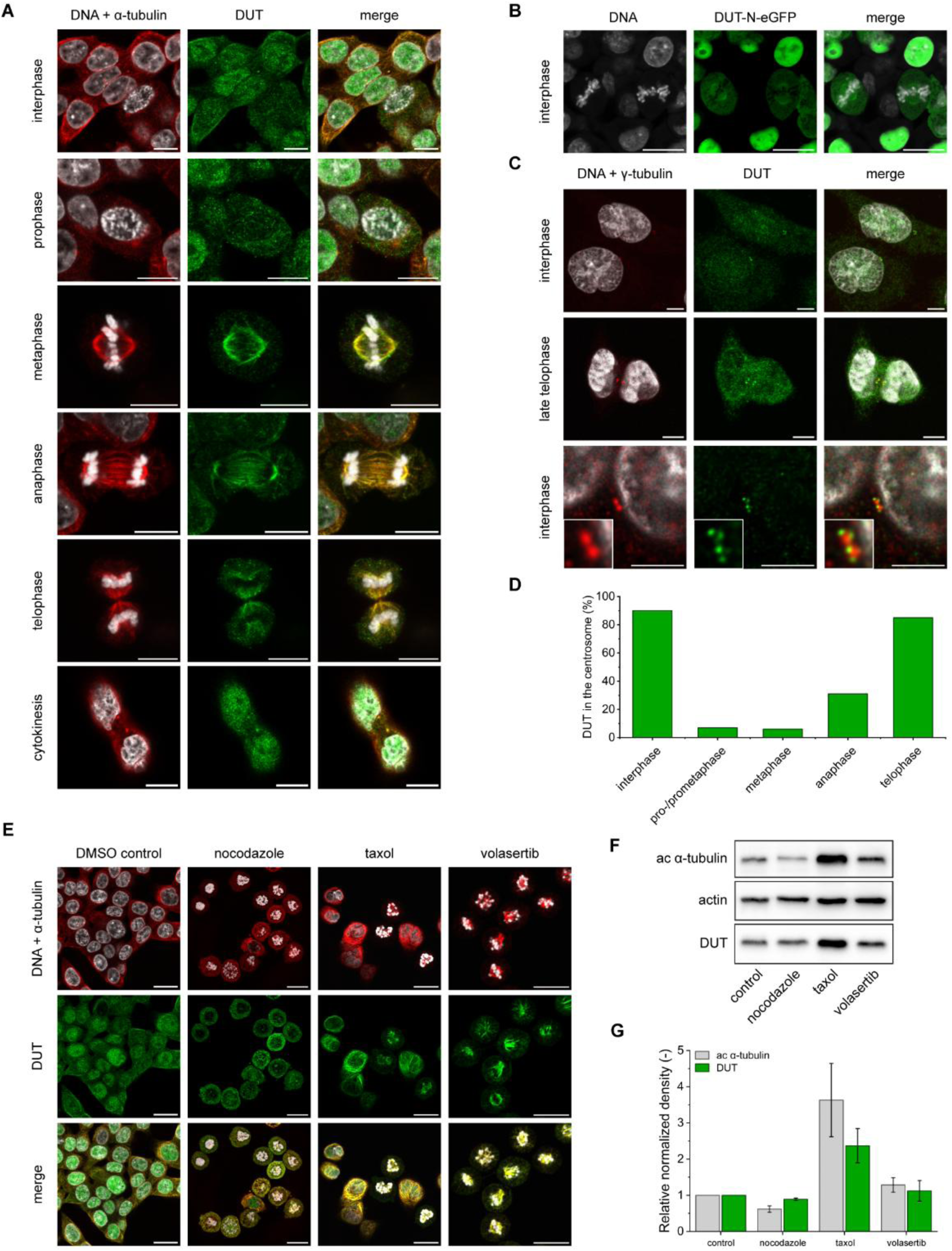
Localization of dUTPase in the different stages of mitosis in human cells. (A) Immunostaining of DNA (gray), α-tubulin (red), and dUTPase (green) in interphase and the different stages of mitosis (prophase, metaphase, anaphase, telophase, and cytokinesis) in HCT116 cells. Scale bar 10 μm. (B) DUT-N-eGFP (green) localization in HCT116 cells stably containing a doxycycline-inducible construct. DNA is shown in gray. Scale bar 20 μm. (C) Immunostaining of DNA (gray), γ-tubulin (red), and dUTPase (green). Magnified sections of the centrosomes are shown in frames. Scale bar 5 μm. (D) Frequency of dUTPase occurrence in the centrosomes in different mitotic phases (n=100 in every phase detected). (E) Immunostaining of DNA (gray), α-tubulin (red), and dUTPase (green) in DMSO, nocodazole-, taxol-, and volasertib-treated HCT116 cells. Scale bar 20 μm. (F-G) Western blot analysis and evaluation of acetylated α-tubulin (red) and dUTPase (green) in DMSO, nocodazole-, taxol-, and volasertib-treated HCT116 cells. Whiskers indicate standard deviation.

Interestingly, dUTPase also formed distinct foci around γ-tubulin at the centrosomes, with stage-specific enrichment observed primarily in interphase and telophase (Figure 3C). Analysis across mitotic stages in HCT116 cells by counting 100 cells in each phase revealed centrosomal localization of dUTPase in more than 85% of interphase and telophase cells (Supplementary Figure S3B, Figure 3D). Moreover, centrosomal localization of dUTPase was observed in ∼30% of anaphase cells, predominantly in late anaphase. Such precise temporal regulation of centrosomal association suggests that dUTPase may coordinate with microtubule-organizing centers to facilitate mitotic progression.

Although dUTPase has been primarily studied as a dNTP pool-sanitizing enzyme, our data indicate a previously unrecognized, dynamic involvement in mitotic architecture. Supporting this, a large-scale dependency analysis across human cancer cell lines (DepMap) reveals that dUTPase is among the most essential proteins identified in cancer, exhibiting extremely strong dependency-predicting features beyond mutation or expression. Moreover, it displays a significant co-dependency with inner centromere protein (INCENP) [42], a core component of the chromosomal passenger complex (CPC). INCENP transitions from broad chromosomal localization to centromeres in early mitosis, to the midzone during anaphase, and ultimately to the midbody in telophase. Furthermore, CDK5 regulatory subunit-associated protein 3 (CDK5RAP3) was identified as one of the top 5 genes, showing co-dependency with dUTPase upon an RNAi screen, according to the DepMap database [43]. Interestingly, CDK5RAP3, which is involved in cell proliferation, cytoskeletal remodeling, migration, and metastasis [44], [45] exhibits similar dynamic re-localization from the centrosomes to the mitotic and central spindle [46]. The close correspondence between the spatial dynamics of dUTPase and these regulatory proteins suggests a multifaceted role that extends beyond nucleotide metabolism. Moreover, bioinformatic analyses of mitotic regulators have independently identified dUTPase among genes co-expressed or physically associated with key centromere/kinetochore components, suggesting a potential mechanistic link [47]. Together, these findings define dUTPase as a dynamically regulated mitotic protein with distinct localization patterns corresponding to specific cell-cycle transitions.

To gain insights into the microtubule-associated localization pattern of dUTPase observed during different mitotic phases, we applied various inhibitors that perturb microtubule dynamics and mitotic processes. We used nocodazole as a microtubule depolymerizing agent and taxol as a microtubule stabilizing agent, respectively, and volasertib to inhibit PLK1 activity in mitosis (Figure 3E). In DMSO control cells, the microtubule network remained intact, as indicated by its filamentous distribution and the presence of asynchronously dividing cells. Upon nocodazole treatment, microtubule assembly is inhibited as indicated by diffuse localization of α-tubulin. In some regions, nucleation of mitotic spindles can be observed, where dUTPase accumulation can also appear, indicating that its localization to the mitotic spindle is microtubule-dependent. Taxol inhibits microtubule depolymerization, thereby stabilizing the microtubule network and increasing acetylated α-tubulin levels in cells [48]. Following taxol treatment, dUTPase exhibits stronger localization to the microtubule network, suggesting its association with acetylated microtubules, and that microtubule dynamics are not required for dUTPase localization to the mitotic spindle. Upon volasertib treatment, cells are synchronized in mitosis, and dUTPase localizes to the mitotic spindle, indicating that PLK1 activity is not required for the observed localization pattern. To determine the protein levels of acetylated α-tubulin and dUTPase following different treatments, we performed Western blot analysis (Figure 3F). Although nocodazole treatment decreased the level of acetylated α-tubulin, dUTPase was not altered; however, upon taxol treatment, its level was markedly increased, while upon volasertib treatment, it was moderately increased, following the acetylated α-tubulin levels. These results suggest that dUTPase levels follow the increase in acetylated α-tubulin (Figure 3G).

### Knockdown of dUTPase in human cells

Several studies have investigated the effect of dUTPase silencing, with a primary focus on sensitivity to fluorouracil derivatives [10], [27], [49], [50]. However, the consequences of sole dUTPase depletion were not investigated in detail; only cell proliferation and dUTP levels were measured [26]. Therefore, we generated an HCT116 cell line stably expressing a cumate-inducible PB-CuO-shDUT cassette for shRNA-mediated silencing. The cassette contains three shRNAs targeting the 3’-UTR of dUTPase and an mOrange reporter (Figure 4A). To determine silencing efficiency, we performed Western blot analysis on dUTPase-silenced (shDUT) HCT116 cells after induction with 100 μg/ml cumate for 48, 72, 96, and 120 hours. As control, we used non-induced cells containing the inducible gene cassette (Supplementary Figure 4A). Following 96-hour cumate induction, the protein level of dUTPase was reduced by ∼95%, indicating effective silencing and near-complete depletion (Figure 4B). Although non-induced control cells have somewhat decreased dUTPase levels (possibly due to leaky expression), no phenotypic changes were observed in our experimental setup, including cell proliferation, DNA damage, cell cycle phase distribution, mitotic defects, or migration rate. Immunostaining revealed efficient depletion of dUTPase in shDUT cells compared to WT and control cells (Figure 4C).

**Figure 4.**
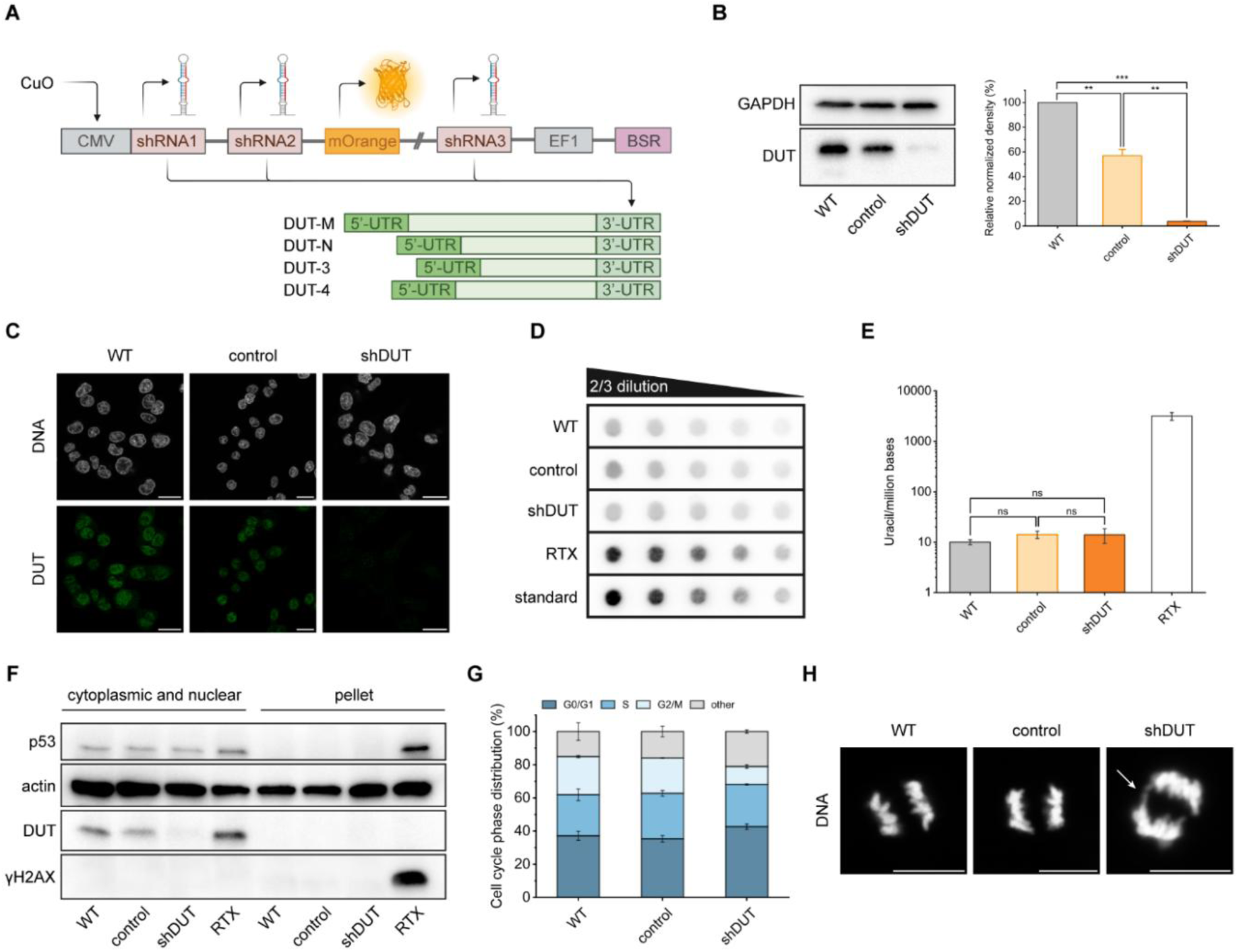
Knockdown of dUTPase in human cells. (A) Schematic representation of the inducible PB-CuO-shDUT construct. Silencing cassette contains cumate-inducible (CuO) CMV promoter (CMV), three shRNA sequences targeting the 3’-UTR of all dUTPase isoforms (shRNA 1, 2, and 3), mOrange fluorescent protein, and blasticidin resistance (BSR) driven by EF1 promoter. (B) Western blot analysis of WT (gray), control (light orange), and shDUT (orange) HCT116 cells. Whiskers indicate standard deviation. Statistical analysis was carried out using One-way ANOVA with Tukey multiple comparisons test (**: p<0.004, ***: p<0.0004). (C) Immunostaining of WT, control, and shDUT HCT116 cells for DNA (gray) and dUTPase (green). Scale bar 20 μm. (D-E) Analysis of genomic uracil level with dot blot in WT, control, shDUT, and RTX-treated HCT116 cells. Whiskers indicate standard deviation. Statistical analysis was carried out using One-way ANOVA with Tukey multiple comparisons test (ns: non-significant). (F) Western blot analysis of γ-H2AX and p53 in cytoplasmic and nuclear and pellet fractions of WT, control, shDUT, and RTX-treated HCT116 cells. (G) Flow cytometry analysis of the cell cycle in WT, control, and shDUT HCT116 cells. n=3 biological replicates. Whiskers indicate standard deviation. (H) DNA staining of WT, control, and shDUT HCT116 anaphase cells. Scale bar 20 μm. Arrow indicates a chromosome bridge.

To test the effect of dUTPase overexpression, we generated an HCT116 cell line stably expressing a doxycycline-inducible PB-TO-DUT-N-eGFP cassette, containing the nuclear isoform of dUTPase with a C-terminal eGFP-tag and an mTagBFP2 reporter (Supplementary Figure 4B). Western blot analysis revealed that dUTPase protein levels increased eightfold compared to WT and non-induced control HCT116 cells (Supplementary Figure 4C).

To test whether genomic uracil levels rise upon dUTPase depletion, we used a dot blot assay [51]. As a positive control, we used raltitrexed-treated (RTX) HCT116 cells stably expressing UGI, a UNG inhibitor. RTX is a clinically relevant, specific thymidylate synthase-targeting chemotherapeutic drug that results in a significantly increased genomic uracil level in UGI-expressing HCT116 cells [52]. To quantify the number of genomic uracils per million bases, we applied a CJ236 *E.* coli strain deficient in both *ung* and *dut*, which has an elevated genomic uracil level [53]. Interestingly, upon dUTPase depletion, we observed no significant change in genomic uracil levels compared to WT and non-induced control cells (Figure 4D and E). It was previously shown that genomic uracil level is not elevated in *dut-*, only in *dut- ung- E.coli* mutants [33], suggesting that sole depletion of dUTPase does not increase genomic uracil level if uracil repair is functional.

To confirm that dUTPase depletion does not induce DNA damage and corresponding repair mechanisms, we determined the levels of γ-H2AX and p53 using Western blot (Figure 4F). The γ-H2AX and p53 signals did not elevate upon dUTPase silencing, indicating that no double-strand breaks are generated and no overactivation of repair mechanisms occurs.

Furthermore, we measured cell cycle phase distribution using flow cytometry and found that dUTPase silencing did not markedly change the cell cycle profile; however, it resulted in a decreased proportion of cells in the G2/M phase (Figure 4G). Notably, cells in S phase did not change upon dUTPase silencing, although the canonical function of dUTPase (dUTP hydrolyzation) is an S-phase-associated function.

Importantly, upon dUTPase depletion, chromosome bridges can be observed in anaphase cells compared to WT and control cells (Figure 4H). Mitotic defects were also observed in *Drosophila* embryos upon silencing both zygotic and maternal dUTPase, highlighting its potential in mitotic processes.

### dUTPase knockdown induces centrosome amplification

Having determined that dUTPase colocalizes with centrosomes in interphase and telophase and loss of dUTPase causes chromosome bridges, it was of immediate interest to check the cellular effects of dUTPase knockdown on centrosome integrity during the cell cycle. Towards this end, we analyzed in depth all phases distinguishable along mitosis, from interphase through pro-/prometaphase, metaphase, anaphase, and telophase (Figure 5). In all phases investigated, 100 cells were analyzed for WT, control, and shDUT cell lines, providing a highly reliable dataset for deciphering physiological effects. We found that dUTPase silencing induced a significant increase in the number of centrosomes, as indicated by the number of γ-tubulin foci, in all phases compared to WT and control cells. Strikingly, a surplus of centrosomes was observed upon dUTPase silencing, and the normal localization pattern of centrosomes was also drastically perturbed (Figure 5).

**Figure 5.**
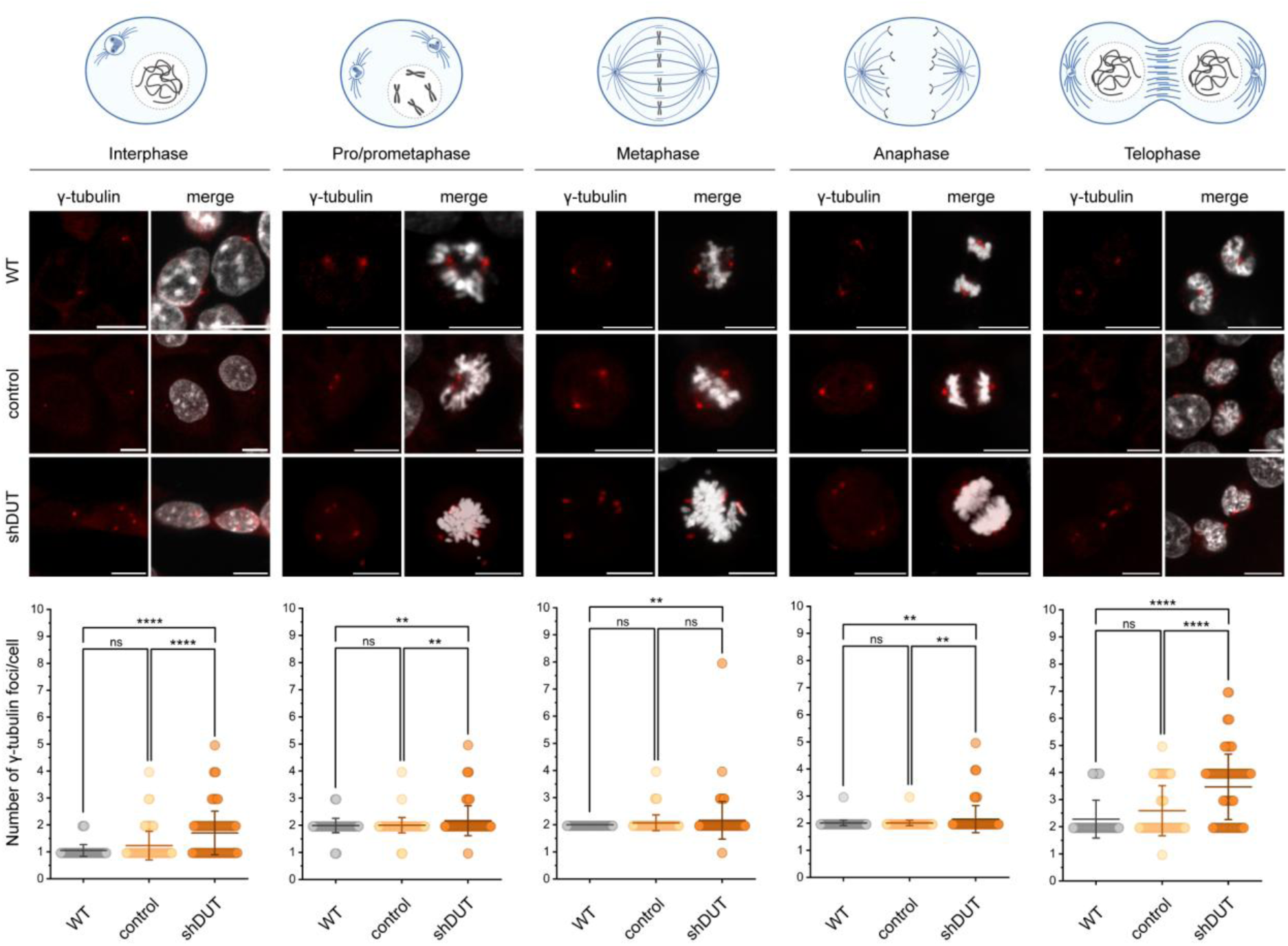
dUTPase knockdown induces centrosome amplification. Representative microscopy images of γ-tubulin (red) and DNA (gray) in WT, control, and shDUT cells in the different cell cycle phases. Scale bar 20 μm. Graphs show the number of γ-tubulin foci per cell in the different cell cycle phases in WT (gray), control (light orange), and shDUT (orange) HCT116 cells. n=100 cells counted in each group of every phase. Whiskers indicate standard deviation. Statistical analysis was carried out using Kruskal-Wallis test with Dunn’s multiple comparisons test (**: p<0.01, ****: p<0.0001, ns: non-significant).

### Effect of dUTPase expression on cell migration

Centrosome integrity and microtubule organization are central not only to accurate mitosis but also to cell polarity and directional migration [54], [55]. Accordingly, centrosome amplification and spindle-associated defects have been linked to altered migratory behavior and metastatic potential through disrupted cytoskeletal dynamics. Given the dynamic association of dUTPase with the mitotic spindle and centrosomes, and the centrosomal defects observed upon its depletion, we next examined whether dUTPase expression levels influence cell migration. Previously, it was shown that RNAi-mediated depletion of dUTPase in HeLa, HT29, and SW620 cells reduces proliferation [26]. To address this, we examined the effects of both dUTPase silencing and overexpression on cell proliferation by determining cell concentration using flow cytometry (Figure 6A and B). Consistent with earlier findings, dUTPase depletion significantly decreased, whereas its overexpression did not affect the proliferation rate.

**Figure 6.**
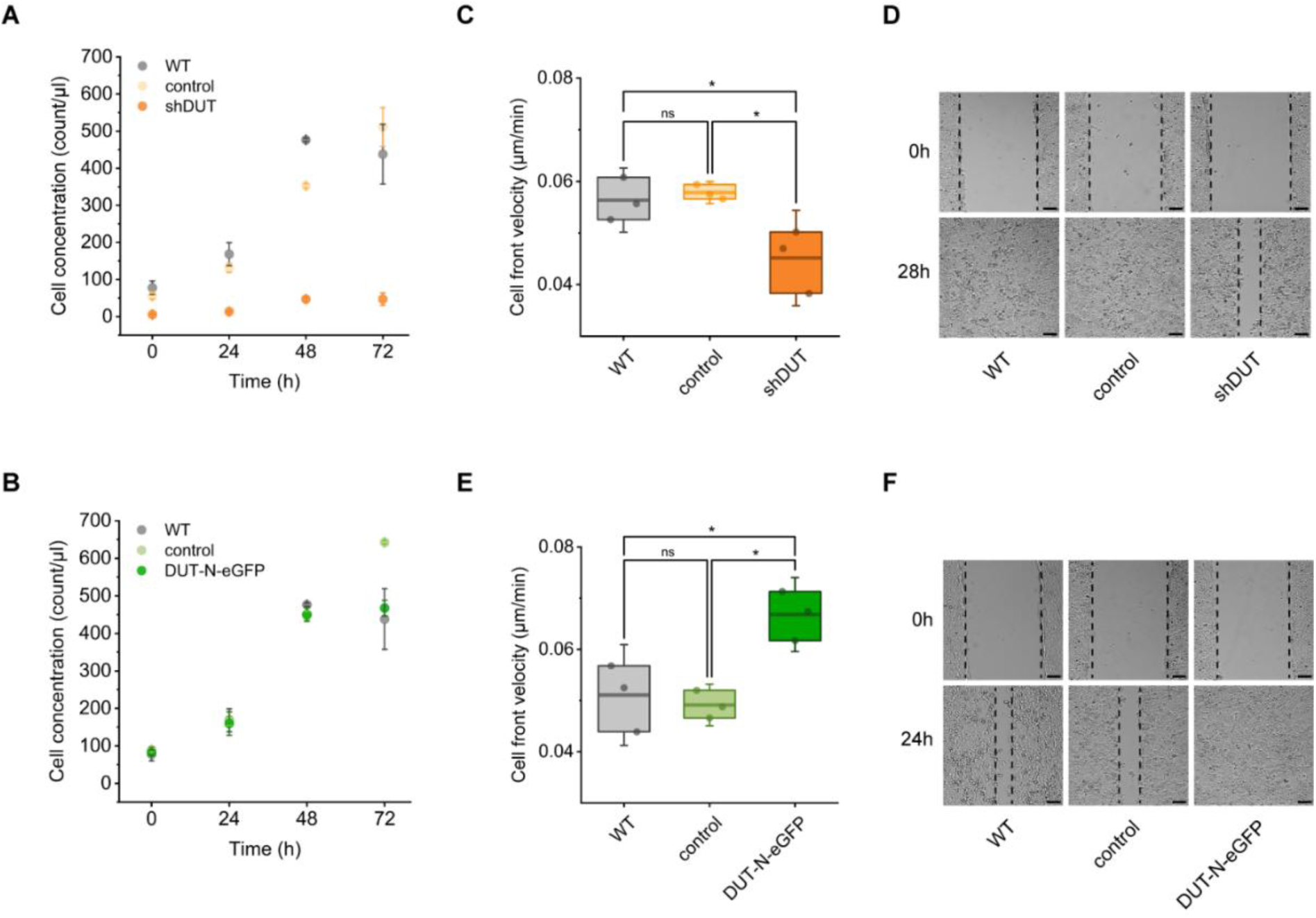
Effect of dUTPase expression on cell migration. (A-B) Cell concentration of WT, control, and (A) shDUT or (B) DUT-N-eGFP overexpressed HCT116 cells. Whiskers indicate standard deviation. (C) Cell front velocity of WT (gray), control (light orange), and shDUT (orange) HCT116 cells. n=3 in each group. Whiskers indicate standard deviation. Statistical analysis was carried out using One-way ANOVA with Tukey multiple comparisons test (*: p<0.05, ns: non-significant). (D) Representative microscopy images of WT, control, and shDUT HCT116 cells at 0h and 28h. Dashed lines indicate cell fronts. Scale bar 100 μm. (E) Cell front velocity of WT (gray), control (light green), and DUT-N-eGFP (green) HCT116 cells. n=3 in each group. Whiskers indicate standard deviation. Statistical analysis was carried out using One-way ANOVA with Tukey multiple comparisons test (*: p <0.05, ns: non-significant). (F) Representative microscopy images of WT, control, and DUT-N-eGFP HCT116 cells at 0h and 24h. Dashed lines indicate cell fronts. Scale bar 100 μm.

Furthermore, we performed a wound healing assay and found that dUTPase depletion significantly reduced (Figure 6C and D), whereas its overexpression significantly increased cell front velocity (Figure 6E and F), relative to WT and non-induced control cells.

Given that the generation time of HCT116 cells exceeds 20 hours, cell division cannot account for the rapid closure of the gap within the less than 30-hour time frame of the experiment, indicating that the assay predominantly measures cell migration rather than proliferation. Although proliferation of shDUT cells was highly reduced, these cells still migrated, albeit at a markedly lower rate. Importantly, dUTPase overexpression did not affect cell proliferation, but substantially increased migration rate. These results clearly demonstrate that dUTPase plays a crucial regulatory role in cell migration. It was previously demonstrated that during mouse neurogenesis, protein expression of dUTPase is highly elevated in the migrating neuroblasts of the rostral migratory stream [7]. Moreover, dUTPase was identified as a reliable biomarker for cancer metastasis [56], [57]. Results of these studies, as well as our current observations, suggest a pioneering function of dUTPase in cellular behaviors relevant to cell migration and metastasis, with further implications for understanding the basis of its role in tissue morphogenesis and metastatic progression.

## Conclusions

The enzymatic activity of dUTPase, which hydrolyzes dUTP to prevent its misincorporation into DNA, has been extensively characterized in the literature [1], [10], [58], [59]. Although several studies [7], [29] and recent reviews [4], [60] have suggested additional, non-canonical functions for dUTPase, to date, no direct experimental evidence has supported such a conception. Here, we identify dUTPase as a mitotic factor and demonstrate that it is indispensable for early embryonic development from *Drosophila* to mouse. Our findings reveal that dUTPase may exert a regulatory function to maintain mitotic and centrosomal integrity.

We showed that in *Drosophila melanogaster*, where the *ung* gene is not encoded in the genome, CRISPR knockout of dUTPase resulted in delayed pupariation and lethality. As maternal dUTPase is present in knockout embryos and dUTPase has low mRNA and protein expression during the larval stage [5], [34], animals can develop until the pupal stage, but not further. In contrast, RNAi-mediated silencing of maternal dUTPase revealed its essential significance during early development, since loss of dUTPase resulted in mitotic defects and erroneous cell division leading to embryonic lethality.

In mouse, knockout of *Dut* was previously demonstrated to cause early embryonic lethality [18], and here we showed that simultaneous knockout of key enzymes involved in uracil DNA repair, Ung and Smug1, failed to rescue *Dut* knockout and likewise resulted in embryonic lethality. In wild-type embryos, we found that dUTPase localized to the mitotic apparatus and showed colocalization with α-tubulin in trophectoderm cells, whereas it accumulated predominantly in the nuclei of the inner cell mass.

Detailed analysis of dUTPase localization throughout mitosis in mammalian cells (HCT116, HEK293, U2OS, MEF) revealed a dynamic spatiotemporal pattern. During interphase, dUTPase is enriched in the nucleus but is also detectable in the close vicinity of centrosomes. Upon entry into mitosis, dUTPase is excluded from the nucleus during prophase, associated with the mitotic spindle during prometaphase, metaphase, and anaphase, and subsequently accumulated at the central spindle and centrosomes during telophase. Moreover, the protein level of dUTPase closely follows changes in acetylated α-tubulin upon taxol treatment, further highlighting its involvement in microtubule-dependent processes.

Previous studies of dUTPase depletion have primarily focused on its role in *de novo* thymidylate metabolism and in the response to chemotherapeutic agents [26], [27], [49], [50]. Here, by specifically investigating mitotic processes, we found that efficient dUTPase knockdown caused chromosome bridges and centrosome amplification, accompanied by a reduction in the G2/M phase cell cycle population and a concomitant decrease in proliferation rate.

The co-dependency between dUTPase and centromere/kinetochore components [47], as well as mitotic regulators, such as INCENP [42] and CDK5RAP3 [43], reported in large-scale, non–hypothesis-driven exploratory studies, independently supports our findings. Moreover, dUTPase depletion decreased the cell migration rate, whereas its overexpression increased it. This phenomenon strongly highlights the biomarker potential of dUTPase in cancer metastasis [56], [57].

Together, our findings redefine dUTPase as an essential mitotic factor with a conserved role in embryonic development. Beyond its well-established enzymatic function in nucleotide metabolism, here we show that dUTPase dynamically associates with the mitotic apparatus and centrosomes to ensure mitotic fidelity, centrosome integrity, and proper cell division. The fact that uracil DNA repair deficiency fails to rescue *Dut* loss *in vivo* suggests that its mitotic function is independent of canonical DNA repair pathways. By linking dUTPase to mitotic spindle dynamics, centrosome homeostasis, and cell migration, our study recognizes dUTPase as a key mitotic factor that contributes to cellular plasticity and developmental processes.

## Materials and Methods

### Symbols used in this study

The following symbols were used to indicate genes and proteins in the different organisms used in this study: *dut* (gene in *Drosophila melanogaster*), Dut (protein in *Drosophila melanogaster*), *Dut* (gene in mouse), *DUT* (gene in human), and DUT (protein in mouse and human).

### *Drosophila* maintenance

*Drosophila* animals were maintained in the HUN-REN Biological Research Centre, Institute of Genetics. *Drosophila* lines used for the genotypes presented: P{MTD-GAL4} (FlyBase ID: FBst0007063), P{His2Av-EGFP.C} (FlyBase ID: FBst0024163), y M{vas-int} w; M{3xP3-RFP.attP’}58A (FlyBase ID: FBst0024484), y M{vas-int.Dm} w; M{1 loxP, attP}51D (kindly gift from Tamás Lukasovics), y v P{nos-phiC31}; attP40 (FlyBase ID: FBst0025709), y v P{nos-phiC31}; attP2 (FlyBase ID: FBst0025710).

### Knockout of *dut* by CRISPR-Cas9 in Drosophila

20 nt long, Cas9 target sequences against dut genomic locus were 5’-GAAAAGGCTGCCGAACCGGA-3’ (site1) and 5’-GCTTTGGCGGATCCCCTCAC-3’ (site2). The gRNA sequences were cloned into the pU6-BbsI-chiRNA plasmid and injected into *Drosophila* embryos with the phsp70-Cas9 plasmid, followed by the flyCRISPR protocol (http://flycrispr.molbio.wisc.edu/) [61]. New mutations were screened for lethality, and deletions were identified by PCR with the following forward and reverse primers (fw: 5’-TATCAGGGTTGGTTGTGATTG-3’ and rev: 5’-ACTCTCACTGCAAGTTGGTTG-3’). Sequence analysis was performed by Sanger sequencing.

### Generation of transgenic rescue constructs in *Drosophila*

To generate rescue constructs, we cloned the upstream 912 bp considered as promoter, and either the Dut-N or Dut-C sequence with a C-terminal 3x Flag epitope into the pUASTattB vector between NheI and XhoI sites, exchanging the UAS regulator for the endogenous promoter. Transgenes were introduced into the *Drosophila* genome by φC31 integrase-mediated transgenesis at attP sites in 58A and 51D genomic regions (2nd chromosome) [62]. Dut-N transgene contains the full-length 23kDa, while Dut-C transgene encodes the shorter 21kDa, NLS-free isoform of dUTPase (dUTPase-PB, FlyBase ID: FBpp0079745).

### Pupariation dynamics of *dut^-/-^* and control animals

Descendant embryos from *dut-*/CyO Act-GFP adults were placed in vials with standard *Drosophila* food. The ratio of puparated *dut^-/-^*homozygote knockout (GFP negative) and *dut-*/CyO Act-GFP heterozygote (used as control, GFP positive) animals was determined from day 5-13 after egg laying.

### Generation of *Drosophila dut* RNAi lines

UAS-driven shRNA against *dut* gene was generated according to the FlyRNAi protocol (https://fgr.hms.harvard.edu/cloning-and-sequencing). A shRNA was designed against the 5’-CTTCATGGTCACTATCAAAGA-3’ of the 3’UTR sequence of dUTPase and cloned into the Valium22 vector [63]. The shRNA transgene was introduced into the *Drosophila* genome by φC31 integrase-mediated transgenesis at attP40 (2nd chromosome) and attP2 (3rd chromosome) sites.

### Maternal depletion of dUTPase and rescue of the phenotypes in *Drosophila*

To knockdown dUTPase in *Drosophila* embryos, we investigated the descendants of P{UAS-*dut* RNAi}attP40/+; P{MTD-GAL4}/{UAS-*dut* RNAi}attP2 females. In the rescue experiments, descendants of P{UAS-*dut* RNAi}attP40/P{Dut}; P{MTD-GAL4}/{UAS-*dut* RNAi}attP2 were investigated. The ratio of laid and hatched eggs, the ratio of hatched and pupating larvae, and the ratio of pupating and adult animals were determined. Experiments were performed at 25°C and 29°C.

### Video microscopy and DAPI staining of *Drosophila* embryos

Embryos laid by P{UAS-*dut* RNAi}attP40/P{His2Av-EGFP.C}; P{MTD-GAL4}/{UAS-*dut* RNAi}attP2 or P{UAS-*dut* RNAi}attP40/P{His2Av-EGFP.C}; +/UAS-*dut* RNAi}attP2 (as control) females were collected 0-1h after egg laying, dechorionated with 50% bleach (2% NaOCl) and washed with water, then aligned on glue glass coverslips and covered with halocarbon oil. Video microscopy was performed using an Olympus CELL-R microscope with a 10x dry objective (NA: 0.3) at a resolution of 0.8 frames per min in time. The embryos between nuclear cycle 9 and 13 (NC9-NC13, 1-2.5h old) were examined. The ratio of damaged nuclei was determined in each embryo in 2 or 3 non-overlapping areas measuring 200x200 pixels (4117 μm2).

For DAPI staining, the embryos laid by P{UAS-*dut* RNAi}attP40/+ P{MTD-GAL4}/{UAS-*dut* RNAi}attP2 or P{UAS-*dut* RNAi}attP40/+; UAS-*dut* RNAi}attP2/+ (as control) females were collected 0-2.5h after egg laying. Embryos were dechorionated with 50% bleach, fixed in 1:1 mixture of heptane:PBS and 4% PFA for 20 min, devitellinized with 1:1 heptane:methanol, rehydrated by washing with 2:1, 1:1, and 1:2 mixture of methanol:PBT (0.1% Triton X-100 in 1×PBS) for 3×5 min, then washed in PBT for 3×10 min. Embryos were stained with 1 μg/ml DAPI in PBT for 30 min, washed with PBT for 3×10 min, and placed on a microscope slide, mounted with Fluoromount-G (Invitrogen). Microscopy was performed using a Leica SP5 AOBS confocal laser scanning microscope with a 20× dry (NA: 0.7) objective.

### Immunohistochemistry of *Drosophila* ovaries

Ovaries of *dut^-/-^*;P{Dut-C} and *dut^-/-^*;P{Dut-N} adult females were dissected, fixed with 4% PFA in PBS for 20 min and washed with PBT for 3×20 min, blocked with PBT-N (1% BSA and 5% FCS in PBT) and incubated overnight with anti-Vasa (rat monoclonal, DSHB, 1:300) and anti-Flag M2 (mouse monoclonal, Sigma F1804, 1:1000) in PBT-N, washed with PBT for 3x20 min, incubated with secondary antibody (Alexa488 conjugated anti-rat, InvitrogenA11006, 1:600 and Alexa647 conjugated anti-mouse Invitrogen A21236, 1:600) and DAPI (1 μg/ml) in PBT-N. Ovaries washed with PBT for 3x20 min were placed on a microscope slide and mounted with Fluoromount-G (Invitrogen). Microscopy was performed using a Leica SP5 AOBS confocal laser scanning microscope with a 63× oil immersion (NA: 1.2) objective.

### Mouse maintenance

Mice were maintained in the Animal Care Facility of the Department of Animal Biotechnology, Institute of Genetics and Biotechnology, Hungarian University of Agriculture and Life Sciences. The present study was conducted in full compliance with the Directive 2010/63/EU, Hungarian Code of Practice for the Care and Use of Animals for Scientific Purposes, and the ARRIVE Guidelines, and in strict accordance with the recommendations and regulations of the European Animal Research Association (https://www.eara.eu/animal-research-law) and the Science Ethics Code of the Hungarian Academy of Sciences (https://mta.hu/data/dokumentumok/english/background/Science_Ethics_Code_English.pdf), including all requirements related to animal welfare and handling; and were approved by the Animal Care and Ethics Committee of the NAIK Agricultural Biotechnology Institute (legal predecessor of the host institution) and the Pest County governmental office (permission number: PEI/001/329-4/2013). For euthanasia, cervical dislocation was performed, and all efforts were made to minimize animal suffering.

FVB/N and CD1 mice were maintained in groups with free access to food and water and were kept under standard conditions, with a light-dark cycle (06:00-18:00 hours) at 22°C. For immunostaining of mouse embryos, 16-cell, 3.5 dpc, and 4.5 dpc embryos were flushed out of the fallopian tube of the fertilized female CD1 mice.

### Gene knockout in mice

*Ung^-/-^ Smug1^-/-^* mice (C57BL/6J background) were kindly provided by Hilde Nilsen [3]. Knock-out of *Dut* was performed by CRISPR/Cas9 as described previously in FVB/N mice [18]. *Ung^-/-^ Smug1^-/-^ Dut^-/-^* embryos were generated by crossing *Ung^-/-^ Smug1^-/-^ Dut^+/-^* mice. Genotyping of offspring was performed by PCR-based analysis of the *Ung, Smug1,* and *Dut* alleles. Genomic DNA was isolated from the tissue samples using a standard phenol-chloroform extraction method. PCR amplification was carried out using MyTaq Red Mix (Meridian Bioscience) according to the manufacturer’s instructions. PCR amplification was carried out in a thermal cycler (ProFlex, Applied Biosystems) with an initial denaturation at 98°C for 5 min, followed by 35 cycles of 98°C for 10 s, 62°C for 25 s, and 72°C for 15 s, and a final extension at 72°C for 5 min. Amplification products were analyzed by agarose gel electrophoresis. Expected banding patterns allowed discrimination of genotypes at the loci of interest. The sequences of the genotyping primers used were as follows: Dut-fw (5’-GGTCGGTGCCTCCTCTAG-3’), Dut-rev (5’-AATAAGCCTTGCACATCCGG-3’), Smug1-WT-fw (5’-GGATGAGGGTTCAGCCAGACCTACA-3’), Smug1-KO-fw (5’-TGACAGGGTCACATGTCGTACATAA-3’), Smug1-WT-rev (5’-ACTGCGAATATGACTTCAGACATCCCG-3’), Smug1-KO-rev (5’-ACTGCGAATATGACTTCAGACATCCC-3’), Ung-WT-fw (5’-CACGGACCTAATCAAGCTCACG-3’), Ung-KO-fw (5’-CTTGGGTGGAGAGGCTATTC-3’), Ung-rev (5’-GGCCCACCCTGACAAATCCCC-3’).

### Genotyping of knockout mouse embryos

3.5 dpc embryos were collected by flushing the fallopian tube of the mother, embedded in gelatin, and fixed 2 days later with 4% PFA for 2 hours at 37°C. Embryos were stained with DAPI at a 1:10.000 dilution (Sigma, D9542) for 15 min at room temperature and analyzed using a Zeiss LSM 710 confocal fluorescence microscope. Genotyping of the embryos was performed after imaging using RT-qPCR. After microscopy analysis, cells were lysed with lysis buffer (10 mM Tris, 1 mM EDTA, 0.2% Triton X-100, pH 8.0) containing 12.5 μg/ml Proteinase K (Sigma, P4850) at 55°C for 1 h, followed by inactivation at 95°C for 15 min. Then, a qPCR followed by melting curve analysis was performed, using the following primers at a final concentration of 400 nM: fw (5’-GGAGATTTTCGGCGGGTAGG-3’) and rev (5’-AATAAGCCTTGCACATCCGG-3’). Amplification was carried out using MyTaq HS Mix (Bioline) and EvaGreen (Biotium, 31000) according to the manufacturer’s instructions. Thermal cycling and detection were carried out in a CFX96 real-time PCR detection system (BioRad) with an initial denaturation at 95°C for 5 min, followed by 30 cycles of 95°C for 30 s, 66°C for 30 s, and 72°C for 30 s, and a final extension at 72°C for 5 min. Melting curve analysis was performed from 66°C to 95°C with an increment of 0.3°C/s, and data were collected every 5 s.

### Immunostaining of mouse embryos

For immunostaining, 16-cell and 3.5 dpc embryos were collected by flushing the mother’s fallopian tube and then transferred to a 24-well plate. They were immediately fixed with PFA for 2 hours at 37°C. For the 6.5 dpc embryos, 3.5 dpc blastocysts were collected with the same method, embedded into gelatin, and fixed 3 days later with 4% PFA for 2 hours at 37°C. Embryos were permeabilized with 3% BSA in PBS (Sigma, A9418) and 0.1% Triton X-100 in PBS for 1 hour. Then embryos were incubated with primary antibodies diluted in 3% BSA in PBS for 2 hours. Samples were washed 3 times with PBS and incubated with secondary antibodies in 3% BSA in PBS for 1 hour, protected from light. After washing with PBS, DNA was stained with a 1:10,000 dilution of DAPI (Sigma, D9542). Then, embryos were transferred to Grace Bio-Labs Culture Well removable chambered coverglass (Sigma, GBL112358) and mounted in FluorSave (Merck Millipore, 345789) reagent. Samples were analyzed with Zeiss LSM710 or Leica SP8 confocal fluorescence microscopes.

### Cell maintenance

The cell lines used in this study were HCT116 (human colorectal carcinoma), MEF (mouse embryonic fibroblast), HEK293 (human embryonic kidney), and U2OS (human osteosarcoma), which were purchased from ATCC. HCT116, MEF, and HEK293 cells were grown in DMEM high glucose GlutaMAX medium (Gibco, 10566024), U2OS cells were grown in McCoy′s 5A Medium (Gibco, 16600082) supplemented with 10% FBS (Gibco, 10500064) and 1% PS (Gibco, 15140-122). Cells were maintained at 37°C in a humidified incubator (Eppendorf, Galaxy 170R) with 5% CO_2_ atmosphere. To verify the absence of mycoplasma infection, we performed genomic DNA isolation and PCR analysis.

### Immunostaining of mammalian cells

For immunostaining, cells (HCT116, MEF, HEK293, and U2OS) were seeded on µ-Slide 8-well microscopy chamber (Ibidi, 80806). For α-tubulin staining, cells were fixed with 4% PFA in PBS for 2 hours at 37°C, while for γ-tubulin staining, cells were fixed with cold methanol for 15 min at 4°C. Cells were permeabilized with 3% BSA in PBS (Sigma, A9418) and 0.1% Triton X-100 in PBS for 1 hour. After, cells were incubated with primary antibodies diluted in 3% BSA in PBS for 2 hours. Samples were washed 3 times with PBS and incubated with secondary antibodies in 3% BSA in PBS for 1 hour, protected from light. After washing with PBS, DNA was stained with a 1:10,000 dilution of DAPI (Sigma, D9542). Cells were then mounted in FluorSave (Merck Millipore, 345789) reagent and analyzed using a Zeiss LSM710 or Leica SP8 confocal fluorescence microscope.

### Knockdown of dUTPase in HCT116 cells

For shRNA-mediated silencing of dUTPase, three shRNA sequences (5’-TTCCGCAATTGAAGGTTGTATG-3’, 5’-CTTCAAGTGTTTTGGTGTTTTG-3’, 5’-AAGCCTGTATTTAACTCATATG-3’) were designed using Dharmacon Horizon to target the 3’-UTR of all dUTPase isoforms. Targeting shRNA sequences was ordered as a mir-30 precursor in the UBC1, ACTB2, and ACTB4 introns of the AddGene multi-shRNA vector (#12391) [64], with the coding sequence of mOrange fluorescent protein. Silencing DNA cassette was cloned into a modified cumate (CuO) inducible PiggyBac (PB) PB-CuO-CMV-MCS-EF1α-CymR-BSD plasmid (System Biosciences). HCT116 cells were co-transfected with PB-CuO-CMV-MCS-EF1α-CymR-BSD and transposase helper vector pRP-mCherry-CAG-hyPBase (Vectorbuilder) in a 2:1 molar ratio with PEI transfection reagent (MedChemPress, HY-K2014) according to the manufacturer’s recommendation. After 2 h post-transfection, antibiotic selection was performed using 9 μg/ml blasticidin (InvivoGen, ant-bl-05). For induction, 100 μg/ml cumate (Sigma, 268402) was used for 96h in all silencing experiments.

### Overexpression of DUT-N in HCT116 cells

For DUT-N overexpression, the coding sequence of the nuclear DUT isoform with C-terminal eGFP fluorescent tag was cloned into a doxycycline-inducible PiggyBac (PB) PB-TO-hNGN2 plasmid, replacing NGN2 (Addgene #172115) with DUT-N-eGFP. HCT116 cells were co-transfected with PB-TO-DUT-N-eGFP and transposase helper vector pRP-mCherry-CAG-hyPBase in a 2:1 molar ratio with PEI transfection reagent (MedChemPress, HY-K2014) according to the manufacturer’s recommendation. After 24h post-transfection, antibiotic selection was performed using 1 μg/ml puromycin (Sigma, P8833). For induction, 0.03 μg/ml doxycycline (Sigma, D3447) was used for 24-72 hours.

### Flow cytometry analysis

To determine the distribution of cells in each cell cycle phase, we used the Click-iT Plus EdU Alexa Fluor 488 Flow Cytometry Assay Kit (Invitrogen, C10632). Cells were seeded in a TC 6-well plate prior to analysis. Cells were treated with 10 μM EdU for 20 min, then collected, centrifuged for 4 min at 200 g, washed with 3% BSA in PBS, and fixed with 4% PFA diluted in PBS for 15 min. Cells were permeabilized, and the click-it reaction was performed according to the manufacturer’s instructions. For the analysis, an Attune NxT flow cytometer (Thermo Fisher Scientific) was used. The EdU Alexa Fluor 488 signal was detected in the BL1 channel with 488 nm excitation and 530/30 nm emission, while the DRAQ5 DNA signal was detected in the RL1 channel with 633 nm excitation and 670/14 nm emission. For data analysis, Attune NxT software version 3.2.1. was used.

For the cell proliferation assay, cell concentration was determined using flow cytometry. Cells were seeded into a 24-well plate (Corning Costar, 3524) in two biological replicates per time point. Cells were collected after 24, 48, 72, and 96 hours, washed with 1% BSA in PBS, and fixed with 4% PFA in PBS for 15 min at room temperature. Samples were stored in 1% BSA in PBS at 4°C until further processing. Cell concentration was measured on 200 μl of fixed samples using an Attune NxT flow cytometer (Thermo Fisher Scientific), and data analysis was carried out using Attune NxT software version 3.2.1.

### Fluorescence-activated cell sorting

Live-cell sorting was performed using a FACSAria III cell sorter (BD Biosciences). Fluorescence of eGFP was excited with a 488 nm laser and detected using a 530/30 nm band-pass filter. Fluorescence of mOrange was excited at 561 nm and detected using a 610/20 nm band-pass filter, while fluorescence of mTagBFP2 was excited at 405 nm and detected using a 450/40 nm band-pass filter. In shDUT HCT116 cells, cells exhibiting low-intensity mOrange-positive fluorescence were gated and sorted, whereas in DUT-N-eGFP overexpressing HCT116 cells, mTagBFP2-positive cells were gated and collected.

### Western blot analysis

For Western blot analysis, 2.5x10^6^ million cells were collected, and washed with PBS twice, then the lysate was fractionated as follows. Cells were resuspended in 200 μl cytoplasmic extraction buffer (20 mM Tris, 10 mM NaCl, 3 mM MgCl2, 0.5 mM DTT, 0.05% NP-40, protease inhibitor tablets (Roche), pH 7.4) and incubated for 15 min on ice, mixing frequently. Next, cells were centrifuged for 7 min at 3000 g (Eppendorf Centrifuge 5425 R) at 4°C, and the supernatant was transferred to a fresh tube and retained as the cytoplasmic fraction. Pellet was then resuspended in 60 μl nuclear extraction buffer (50 mM Tris, 150 mM NaCl, 50 mM NaF, 5 mM EDTA, 1 mM EGTA, 1 mM PMSF, 1% NP-40, protease inhibitor tablets (Roche), pH 7.4) and incubated on ice for 30 min, vortexing every 5 min. Samples were centrifuged at 16,000 g for 10 min at 4°C, and the supernatant was collected as the nuclear fraction. The pellet was resuspended in 30 mL PBS as the insoluble fraction. Samples were incubated at 95°C in 5X loading buffer (250 mM Tris-HCl, 50% glycerol, 10% DTT, 10% SDS, 0.05% bromophenol blue, pH 6.8) and then loaded onto a 12% polyacrylamide gel. As a protein ladder, GRS Protein Marker MultiColor (GRiSP, GLP01.0500) was used. Electrophoresis was performed in Running Buffer (25 mM Tris–HCl, 20 mM glycine, 0.1% SDS) for 1 hour at 200 V in the Mini-PROTEAN Electrophoresis System (Bio-Rad). The transfer was carried out in transfer buffer (10 mM CAPS, 15% methanol, pH 11) for 3 hours at 250 mA, using Immobilon-P 0.45 µm pore size PVDF transfer membrane (Merck, IPVH09120). After transfer, the membrane was cut and placed in TBS-T (25 mM Tris-HCl, 140 mM NaCl, 3 mM KCl, 0.05% Tween-20, pH 7.4). The membrane was blocked in 3% BSA diluted in TBS-T for 1 hour and then incubated overnight with primary antibodies, anti-actin and anti-dUTPase. The next day, the membrane was incubated with HRP-conjugated secondary antibodies, washed 3 times with TBS-T for 10 min, and then placed in TBS buffer until imaging. Bands were visualized with Immobilon Western Chemiluminescent HRP Substrate (Merck, WBKLS0100). Imaging was performed using a ChemiDoc MP Imaging System (Bio-Rad).

### Wound healing assay

HCT116 cells were seeded on 24-well plate (Corning Costar, 3524) applying removable 2-well cell culture inserts (Ibidi, 80209) using three biological replicates. After 24h incubation inserts were removed, cells were washed with PBS and supplied with fresh media. Plate was transferred to JuLi Stage microscope (NanoEnTek) and data was collected in every 4 hours over time course of 30 hours using a 4x objective. Data analysis was performed using Fiji (https://fiji.sc/) [65] and cell front velocity was determined according to Ibidi’s recommendations.

### Software used in this study

Microscopic images were acquired using Leica LAS X software (Leica Microsystems GmbH, v 3.10.0) or ZEN black edition software (Carl Zeiss Microscopy GmbH, v 14.0.0.201). Post-processing of images, including adjustment and inserting scale bar was performed using Fiji (https://fiji.sc/) [65]. Western blot and dot blot images were processed and analyzed using Image Lab software (BioRad Laboratories, v 6.1.0). Graphs were generated using Origin 2018 (OriginLab Corporation, v b9.5.1.195). Schematic figures were generated with BioRender. Final figures were assembled using Adobe Illustrator CC 2020 (Adobe Inc., v 25.3.1).

## Acknowledgement

This work was supported by the National Research, Development and Innovation Fund of Hungary (K135231, K138318, K146890, FK137867, NKP-2018-1.2.1-NKP-2018-00005, 2022-1.2.2-TÉT-IPARI-UZ-2022-00003), the TKP2021-EGA-02 grant, implemented with support provided by the Ministry for Innovation and Technology of Hungary from the National Research, Development and Innovation Office, and the ICGEB Research Grants Programme 2023 (CRP/HUN23-02). This research was also supported by the National Research, Development and Innovation Office and by the Agribiotechnology and Precision Breeding for Food Security National Laboratory (RRF-2.3.1-21-2022-00007). We thank the Drosophila Injection Service (Institute of Genetics, HUN-REN) for generating transgenic *Drosophila* lines. We are grateful to Kate M. O’Connor-Giles (University of Wisconsin) for providing CRISPR-Cas9 tools used to generate the *dut* mutant in *Drosophila*. We also acknowledge the microscopy support provided by the Cellular Imaging Laboratory, Biological Research Centre, HUN-REN. G.R. is supported by the Momentum Grant of the Hungarian Academy of Sciences (LP2023-15/2023), EMBO Installation Grant (IG5670-2024), and the HUN-REN Welcome Home and Foreign Researcher Recruitment Grant (KSZF-143/2023). This work was partially supported by the Research Council of Norway through its Centres of Excellence scheme, Project Number 33271. We thank Éva Tankó for providing materials for Western blot analysis and molecular cloning. Furthermore, we appreciate András Füredi and Eszter Bajtai for providing materials and expertise in migration experiments. The project was also supported by the Doctoral Excellence Fellowship Programme (DCEP) is funded by the National Research Development and Innovation Fund of the Ministry of Culture and Innovation, and the Budapest University of Technology and Economics.

## Author contributions

B.G.V., N.N., O.T., J.T., G.R., L.H., M.E., G.A.R., and E.G. conceived the projects. L.H. and M.E. provided materials and performed *Drosophila* experiments. N.N., O.T., G.A.R., T.P., O.I.H., L.H., and E.G. provided materials and expertise for mouse experiments and performed the experiments. H.L.N. provided the *Ung^-/-^ Smug1^-/-^* mouse strain. M.U. and E.G. provided materials and expertise for mouse embryo immunostaining. N.N., O.T., and E.O. designed and performed experiments on mammalian cells and analyzed the data. F.B.V. and Z.R.G. assisted with experimental work. G.R. provided advice and materials for silencing and overexpression experiments in human cells. E.S. and G.V. provided expertise in FACS and flow cytometry analysis. J.T., A.B., B.G.V., and G.R. provided professional advice and contributed to the analysis of experimental data. N.N., O.T., E.O., J.T., and B.G.V. wrote the manuscript and created figures. All authors contributed to manuscript editing.

## Supplementary Information

**Supplementary Figure S1. Loss of dUTPase in *Drosophila melanogaster* leads to mitotic defects and lethality.**

**Supplementary Figure S2. Investigation of *Dut* knockout and WT mouse embryos.**

**Supplementary Figure S3. Localization of dUTPase in the different stages of mitosis.**

**Supplementary Figure S4. Knockdown and overexpression of dUTPase in human cells.**

**Supplementary Video S1. Video microscopy of control embryos of *Drosophila melanogaster*.**

**Supplementary Video S2. Video microscopy of dUTPase RNAi embryos of *Drosophila melanogaster*.**

## References

[1] B. G. Vértessy and J. Tóth, “Keeping uracil out of DNA: physiological role, structure and catalytic mechanism of dUTPases.,” Acc. Chem. Res., vol. 42, no. 1, pp. 97–106, Jan. 2009, doi: 10.1021/ar800114w.

[2] L. A. Frederico, T. A. Kunkel, and B. R. Shaw, “A sensitive genetic assay for the detection of cytosine deamination: determination of rate constants and the activation energy.,” Biochemistry, vol. 29, no. 10, pp. 2532–2537, Mar. 1990, doi: 10.1021/bi00462a015.

[3] L. Alsøe et al., “Uracil accumulation and mutagenesis dominated by cytosine deamination in cpg dinucleotides in mice lacking UNG and SMUG1.,” Sci. Rep., vol. 7, no. 1, p. 7199, Aug. 2017, doi: 10.1038/s41598-017-07314-5.

[4] O. Mortusewicz, J. Haslam, H. Gad, and T. Helleday, “Uracil-induced replication stress drives mutations, genome instability, anti-cancer treatment efficacy, and resistance.,” Mol. Cell, vol. 85, no. 10, pp. 1897–1906, May 2025, doi: 10.1016/j.molcel.2025.04.015.

[5] A. Horváth et al., “dUTPase expression correlates with cell division potential in Drosophila melanogaster.,” FEBS J., vol. 282, no. 10, pp. 1998–2013, May 2015, doi: 10.1111/febs.13255.

[6] J. R. Strahler et al., “Maturation stage and proliferation-dependent expression of dUTPase in human T cells.,” Proc Natl Acad Sci USA, vol. 90, no. 11, pp. 4991–4995, Jun. 1993, doi: 10.1073/pnas.90.11.4991.

[7] N. Nagy et al., “Characterization of dUTPase expression in mouse postnatal development and adult neurogenesis.,” Sci. Rep., vol. 14, no. 1, p. 13139, Jun. 2024, doi: 10.1038/s41598-024-63405-0.

[8] G. A. Rácz, N. Nagy, G. Várady, J. Tóvári, Á. Apáti, and B. G. Vértessy, “Discovery of two new isoforms of the human DUT gene.,” Sci. Rep., vol. 13, no. 1, p. 7760, May 2023, doi: 10.1038/s41598-023-32970-1.

[9] R. D. Ladner et al., “dUTP nucleotidohydrolase isoform expression in normal and neoplastic tissues: association with survival and response to 5-fluorouracil in colorectal cancer.,” Cancer Res., vol. 60, no. 13, pp. 3493–3503, Jul. 2000.

[10] R. D. Ladner, “The role of dUTPase and uracil-DNA repair in cancer chemotherapy.,” Curr. Protein Pept. Sci., vol. 2, no. 4, pp. 361–370, Dec. 2001, doi: 10.2174/1389203013380991.

[11] R. D. Ladner, D. E. McNulty, S. A. Carr, G. D. Roberts, and S. J. Caradonna, “Characterization of distinct nuclear and mitochondrial forms of human deoxyuridine triphosphate nucleotidohydrolase.,” J. Biol. Chem., vol. 271, no. 13, pp. 7745–7751, Mar. 1996, doi: 10.1074/jbc.271.13.7745.

[12] R. D. Ladner and S. J. Caradonna, “The human dUTPase gene encodes both nuclear and mitochondrial isoforms. Differential expression of the isoforms and characterization of a cDNA encoding the mitochondrial species.,” J. Biol. Chem., vol. 272, no. 30, pp. 19072–19080, Jul. 1997, doi: 10.1074/jbc.272.30.19072.

[13] R. D. Ladner, S. A. Carr, M. J. Huddleston, D. E. McNulty, and S. J. Caradonna, “Identification of a consensus cyclin-dependent kinase phosphorylation site unique to the nuclear form of human deoxyuridine triphosphate nucleotidohydrolase.,” J. Biol. Chem., vol. 271, no. 13, pp. 7752–7757, Mar. 1996, doi: 10.1074/jbc.271.13.7752.

[14] G. Róna et al., “Phosphorylation adjacent to the nuclear localization signal of human dUTPase abolishes nuclear import: structural and mechanistic insights.,” Acta Crystallogr. D Biol. Crystallogr., vol. 69, no. Pt 12, pp. 2495–2505, Dec. 2013, doi: 10.1107/S0907444913023354.

[15] G. Róna et al., “Dynamics of re-constitution of the human nuclear proteome after cell division is regulated by NLS-adjacent phosphorylation.,” Cell Cycle, vol. 13, no. 22, pp. 3551–3564, 2014, doi: 10.4161/15384101.2014.960740.

[16] M. H. Gadsden, E. M. McIntosh, J. C. Game, P. J. Wilson, and R. H. Haynes, “dUTP pyrophosphatase is an essential enzyme in Saccharomyces cerevisiae.,” EMBO J., vol. 12, no. 11, pp. 4425–4431, Nov. 1993, doi: 10.1002/j.1460-2075.1993.tb06127.x.

[17] H. Kumar et al., “Functional genetic evaluation of DNA house-cleaning enzymes in the malaria parasite: dUTPase and Ap4AH are essential in Plasmodium berghei but ITPase and NDH are dispensable.,” Expert Opin. Ther. Targets, vol. 23, no. 3, pp. 251–261, Mar. 2019, doi: 10.1080/14728222.2019.1575810.

[18] H. L. Pálinkás et al., “CRISPR/Cas9-Mediated Knock-Out of dUTPase in Mice Leads to Early Embryonic Lethality.,” Biomolecules, vol. 9, no. 4, Apr. 2019, doi: 10.3390/biom9040136.

[19] V. Perey-Simon et al., “dUTPase is essential in zebrafish development and possesses several single-nucleotide variants with pronounced structural and functional consequences.,” FEBS Open Bio, Dec. 2025, doi: 10.1002/2211-5463.70176.

[20] N. Siaud et al., “The SOS screen in Arabidopsis: a search for functions involved in DNA metabolism.,” DNA Repair (Amst), vol. 9, no. 5, pp. 567–578, May 2010, doi: 10.1016/j.dnarep.2010.02.009.

[21] E. Dubois et al., “Homologous recombination is stimulated by a decrease in dUTPase in Arabidopsis.,” PLoS ONE, vol. 6, no. 4, p. e18658, Apr. 2011, doi: 10.1371/journal.pone.0018658.

[22] V. M. Castillo-Acosta, A. M. Estévez, A. E. Vidal, L. M. Ruiz-Perez, and D. González-Pacanowska, “Depletion of dimeric all-alpha dUTPase induces DNA strand breaks and impairs cell cycle progression in Trypanosoma brucei.,” Int. J. Biochem. Cell Biol., vol. 40, no. 12, pp. 2901–2913, Jul. 2008, doi: 10.1016/j.biocel.2008.06.009.

[23] M. Guillet, P. A. Van Der Kemp, and S. Boiteux, “dUTPase activity is critical to maintain genetic stability in Saccharomyces cerevisiae.,” Nucleic Acids Res., vol. 34, no. 7, pp. 2056–2066, Apr. 2006, doi: 10.1093/nar/gkl139.

[24] M. Dengg, T. Garcia-Muse, S. G. Gill, N. Ashcroft, S. J. Boulton, and H. Nilsen, “Abrogation of the CLK-2 checkpoint leads to tolerance to base-excision repair intermediates.,” EMBO Rep., vol. 7, no. 10, pp. 1046–1051, Oct. 2006, doi: 10.1038/sj.embor.7400782.

[25] V. Muha et al., “Uracil-containing DNA in Drosophila: stability, stage-specific accumulation, and developmental involvement.,” PLoS Genet., vol. 8, no. 6, p. e1002738, Jun. 2012, doi: 10.1371/journal.pgen.1002738.

[26] A. W. Studebaker, W. P. Lafuse, R. Kloesel, and M. V. Williams, “Modulation of human dUTPase using small interfering RNA.,” Biochem. Biophys. Res. Commun., vol. 327, no. 1, pp. 306–310, Feb. 2005, doi: 10.1016/j.bbrc.2004.12.021.

[27] G. Merényi et al., “Cellular response to efficient dUTPase RNAi silencing in stable HeLa cell lines perturbs expression levels of genes involved in thymidylate metabolism.,” Nucleosides Nucleotides Nucleic Acids, vol. 30, no. 6, pp. 369–390, Jun. 2011, doi: 10.1080/15257770.2011.582849.

[28] A. F. Taylor and B. Weiss, “Role of exonuclease III in the base excision repair of uracil-containing DNA.,” J. Bacteriol., vol. 151, no. 1, pp. 351–357, Jul. 1982, doi: 10.1128/jb.151.1.351-357.1982.

[29] H. H. el-Hajj, H. Zhang, and B. Weiss, “Lethality of a dut (deoxyuridine triphosphatase) mutation in Escherichia coli.,” J. Bacteriol., vol. 170, no. 3, pp. 1069–1075, Mar. 1988, doi: 10.1128/jb.170.3.1069-1075.1988.

[30] E. A. Kouzminova and A. Kuzminov, “Chromosomal fragmentation in dUTPase-deficient mutants of Escherichia coli and its recombinational repair.,” Mol. Microbiol., vol. 51, no. 5, pp. 1279–1295, Mar. 2004, doi: 10.1111/j.1365-2958.2003.03924.x.

[31] B. K. Tye and I. R. Lehman, “Excision repair of uracil incorporated in DNA as a result of a defect in dUTPase.,” J. Mol. Biol., vol. 117, no. 2, pp. 293–306, Dec. 1977, doi: 10.1016/0022-2836(77)90128-0.

[32] S. J. Hochhauser and B. Weiss, “Escherichia coli mutants deficient in deoxyuridine triphosphatase.,” J. Bacteriol., vol. 134, no. 1, pp. 157–166, Apr. 1978, doi: 10.1128/jb.134.1.157-166.1978.

[33] H. R. Warner, B. K. Duncan, C. Garrett, and J. Neuhard, “Synthesis and metabolism of uracil-containing deoxyribonucleic acid in Escherichia coli.,” J. Bacteriol., vol. 145, no. 2, pp. 687–695, Feb. 1981, doi: 10.1128/jb.145.2.687-695.1981.

[34] A. Békési et al., “Developmental regulation of dUTPase in Drosophila melanogaster.,” J. Biol. Chem., vol. 279, no. 21, pp. 22362–22370, May 2004, doi: 10.1074/jbc.M313647200.

[35] M. Carter et al., “Crystal structure, biochemical and cellular activities demonstrate separate functions of MTH1 and MTH2.,” Nat. Commun., vol. 6, p. 7871, Aug. 2015, doi: 10.1038/ncomms8871.

[36] T. Ogawa et al., “Molecular characterization of organelle-type Nudix hydrolases in Arabidopsis.,” Plant Physiol., vol. 148, no. 3, pp. 1412–1424, Nov. 2008, doi: 10.1104/pp.108.128413.

[37] J. Rehwinkel et al., “SAMHD1-dependent retroviral control and escape in mice.,” EMBO J., vol. 32, no. 18, pp. 2454–2462, Sep. 2013, doi: 10.1038/emboj.2013.163.

[38] Z. Shu, S. Row, and W.-M. Deng, “Endoreplication: the good, the bad, and the ugly.,” Trends Cell Biol., vol. 28, no. 6, pp. 465–474, Jun. 2018, doi: 10.1016/j.tcb.2018.02.006.

[39] V. Muha, I. Zagyva, Z. Venkei, J. Szabad, and B. G. Vértessy, “Nuclear localization signal-dependent and -independent movements of Drosophila melanogaster dUTPase isoforms during nuclear cleavage.,” Biochem. Biophys. Res. Commun., vol. 381, no. 2, pp. 271–275, Apr. 2009, doi: 10.1016/j.bbrc.2009.02.036.

[40] A. Courtois and T. Hiiragi, “Gradual meiosis-to-mitosis transition in the early mouse embryo.,” Results Probl. Cell Differ., vol. 55, pp. 107–114, 2012, doi: 10.1007/978-3-642-30406-4_6.

[41] A. Courtois, M. Schuh, J. Ellenberg, and T. Hiiragi, “The transition from meiotic to mitotic spindle assembly is gradual during early mammalian development.,” J. Cell Biol., vol. 198, no. 3, pp. 357–370, Aug. 2012, doi: 10.1083/jcb.201202135.

[42] A. Tsherniak et al., “Defining a cancer dependency map.,” Cell, vol. 170, no. 3, pp. 564–576.e16, Jul. 2017, doi: 10.1016/j.cell.2017.06.010.

[43] R. Arafeh, T. Shibue, J. M. Dempster, W. C. Hahn, and F. Vazquez, “The present and future of the Cancer Dependency Map.,” Nat. Rev. Cancer, vol. 25, no. 1, pp. 59–73, Jan. 2025, doi: 10.1038/s41568-024-00763-x.

[44] G. W.-Y. Mak et al., “Overexpression of a novel activator of PAK4, the CDK5 kinase-associated protein CDK5RAP3, promotes hepatocellular carcinoma metastasis.,” Cancer Res., vol. 71, no. 8, pp. 2949–2958, Apr. 2011, doi: 10.1158/0008-5472.CAN-10-4046.

[45] Y .-F. Dai et al., “LZAP promotes the proliferation and invasiveness of cervical carcinoma cells by targeting AKT and EMT.,” J. Cancer, vol. 11, no. 6, pp. 1625–1633, Jan. 2020, doi: 10.7150/jca.39359.

[46] H. Yan et al., “CDK5RAP3, an essential regulator of checkpoint, interacts with RPL26 and maintains the stability of cell growth.,” Cell Prolif., vol. 55, no. 5, p. e13240, May 2022, doi: 10.1111/cpr.13240.

[47] A. R. Tipton, K. Wang, P. Oladimeji, S. Sufi, Z. Gu, and S.-T. Liu, “Identification of novel mitosis regulators through data mining with human centromere/kinetochore proteins as group queries.,” BMC Cell Biol., vol. 13, p. 15, Jun. 2012, doi: 10.1186/1471-2121-13-15.

[48] S. Cang, Y. Ma, J.-W. Chiao, and D. Liu, “Phenethyl isothiocyanate and paclitaxel synergistically enhanced apoptosis and alpha-tubulin hyperacetylation in breast cancer cells.,” Exp. Hematol. Oncol., vol. 3, no. 1, p. 5, Feb. 2014, doi: 10.1186/2162-3619-3-5.

[49] S. E. Koehler and R. D. Ladner, “Small interfering RNA-mediated suppression of dUTPase sensitizes cancer cell lines to thymidylate synthase inhibition.,” Mol. Pharmacol., vol. 66, no. 3, pp. 620–626, Sep. 2004, doi: 10.1124/mol.66.3.

[50] P. M. Wilson, M. J. LaBonte, H.-J. Lenz, P. C. Mack, and R. D. Ladner, “Inhibition of dUTPase induces synthetic lethality with thymidylate synthase-targeted therapies in non-small cell lung cancer.,” Mol. Cancer Ther., vol. 11, no. 3, pp. 616–628, Mar. 2012, doi: 10.1158/1535-7163.MCT-11-0781.

[51] G. Róna et al., “Detection of uracil within DNA using a sensitive labeling method for in vitro and cellular applications.,” Nucleic Acids Res., vol. 44, no. 3, p. e28, Feb. 2016, doi: 10.1093/nar/gkv977.

[52] H. L. Pálinkás et al., “Genome-wide alterations of uracil distribution patterns in human DNA upon chemotherapeutic treatments.,” eLife, vol. 9, Sep. 2020, doi: 10.7554/eLife.60498.

[53] A. Horváth and B. G. Vértessy, “A one-step method for quantitative determination of uracil in DNA by real-time PCR.,” Nucleic Acids Res., vol. 38, no. 21, p. e196, Nov. 2010, doi: 10.1093/nar/gkq815.

[54] N. M. Wakida, E. L. Botvinick, J. Lin, and M. W. Berns, “An intact centrosome is required for the maintenance of polarization during directional cell migration.,” PLoS ONE, vol. 5, no. 12, p. e15462, Dec. 2010, doi: 10.1371/journal.pone.0015462.

[55] N. Tang and W. F. Marshall, “Centrosome positioning in vertebrate development.,” J. Cell Sci., vol. 125, no. Pt 21, pp. 4951–4961, Nov. 2012, doi: 10.1242/jcs.038083.

[56] J. Fleischmann et al., “Expression of deoxyuridine triphosphatase (dUTPase) in colorectal tumours,” International Journal of Cancer, Dec. 1999., 10.1002/(sici)1097-0215(19991222)84:6<614::aid-ijc13>3.0.co;2-p

[57] A. Kawahara et al., “Higher expression of deoxyuridine triphosphatase (dUTPase) may predict the metastasis potential of colorectal cancer.,” J. Clin. Pathol., vol. 62, no. 4, pp. 364–369, Apr. 2009, doi: 10.1136/jcp.2008.060004.

[58] I. Leveles et al., “Structure and enzymatic mechanism of a moonlighting dUTPase.,” Acta Crystallogr. D Biol. Crystallogr., vol. 69, no. Pt 12, pp. 2298–2308, Dec. 2013, doi: 10.1107/S0907444913021136.

[59] R. Persson, E. S. Cedergren-Zeppezauer, and K. S. Wilson, “Homotrimeric dUTPases; structural solutions for specific recognition and hydrolysis of dUTP.,” Curr. Protein Pept. Sci., vol. 2, no. 4, pp. 287–300, Dec. 2001, doi: 10.2174/1389203013381035.

[60] A. Glukhov, U. Dzhus, I. Kolyadenko, G. Selikhanov, and A. Gabdulkhakov, “Is dUTPase Enzymatic Activity Truly Essential for Viability?,” ijms, vol. 26, no. 19, p. 9260, Sep. 2025, doi: 10.3390/ijms26199260.

[61] S. J. Gratz et al., “Genome engineering of Drosophila with the CRISPR RNA-guided Cas9 nuclease.,” Genetics, vol. 194, no. 4, pp. 1029–1035, Aug. 2013, doi: 10.1534/genetics.113.152710.

[62] J. Bischof, R. K. Maeda, M. Hediger, F. Karch, and K. Basler, “An optimized transgenesis system for Drosophila using germ-line-specific phiC31 integrases.,” Proc Natl Acad Sci USA, vol. 104, no. 9, pp. 3312–3317, Feb. 2007, doi: 10.1073/pnas.0611511104.

[63] J.-Q. Ni et al., “A genome-scale shRNA resource for transgenic RNAi in Drosophila.,” Nat. Methods, vol. 8, no. 5, pp. 405–407, May 2011, doi: 10.1038/nmeth.1592.

[64] X .-G. Xia, H. Zhou, and Z. Xu, “Multiple shRNAs expressed by an inducible pol II promoter can knock down the expression of multiple target genes.,” BioTechniques, vol. 41, no. 1, pp. 64–68, Jul. 2006, doi: 10.2144/000112198.

[65] J. Schindelin et al., “Fiji: an open-source platform for biological-image analysis.,” Nat. Methods, vol. 9, no. 7, pp. 676–682, Jun. 2012, doi: 10.1038/nmeth.2019.

