## Supplementary Information for "Recognizing dUTPase as a mitotic factor essential for early embryonic development"

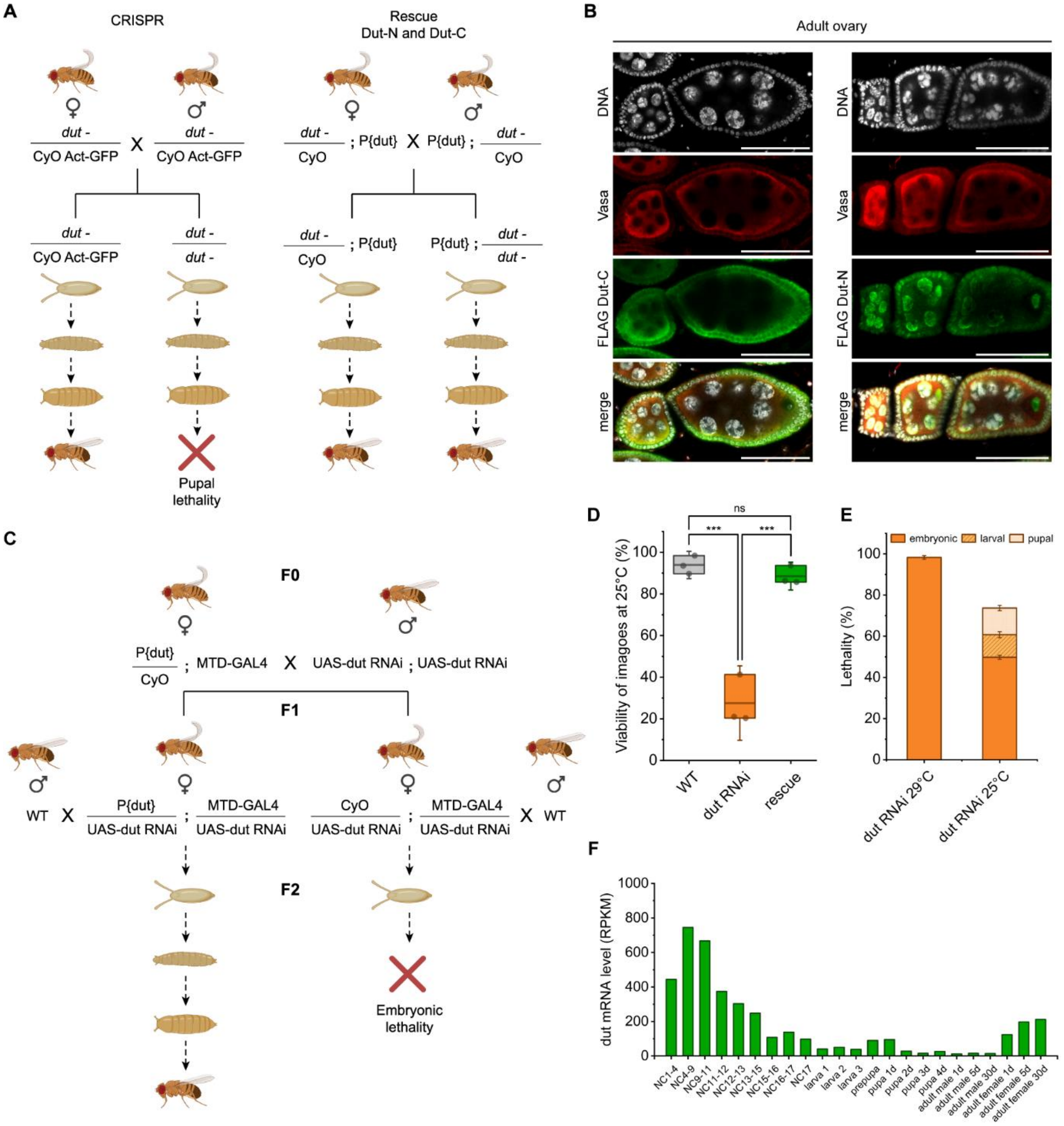


**Supplementary Figure S1. Loss of dUTPase in *Drosophila melanogaster* leads to mitotic defects and lethality.** (A) Schematic representation of *Drosophila melanogaster* crossings to generate CRISPR knockout of *dut,* and rescue with Dut-N or Dut-C transgene with its native promoter (P{dut}). (B) Microscopy images of *Drosophila* ovaries showing DNA (gray), Vasa (red), FLAG-tagged Dut-N or Dut-C rescue constructs (green). Scale bar 50 µm. (C) Schematic representation of crossings of *Drosophila melanogaster* to generate maternally driven silencing (MTD-GAL4/UAS-*dut* RNAi) and/or rescue with Dut-N transgene with their shared native promoter (P{dut}). (D) Viability of dUTPase knockdown animals (orange) at 25°C compared to wild type (WT, gray) and Dut-N rescue (green) imagoes. Whiskers show standard deviation. Statistical analysis was carried out using One-way ANOVA with Tukey multiple comparisons test (***: $p<0.002$, ns: non-significant). (E) Lethality of dUTPase knockdown animals at 29°C and 25°C. Whiskers show standard deviation. (F) mRNA level of dUTPase during *Drosophila* developmental stages according to modENCODE mRNA data available on FlyBase**.**

**Supplementary Video S1. Video microscopy of control embryos of *Drosophila melanogaster.*** Chromosomes in early embryos were visualized by GFP signal expressed from the H2Av-eGFP transgene. Scale bar 50 µm.

**Supplementary Video S2. Video microscopy of dUTPase RNAi embryos of *Drosophila melanogaster*.** Chromosomes in early embryos were visualized by GFP signal expressed from the H2Av-eGFP transgene. Scale bar 50 µm.

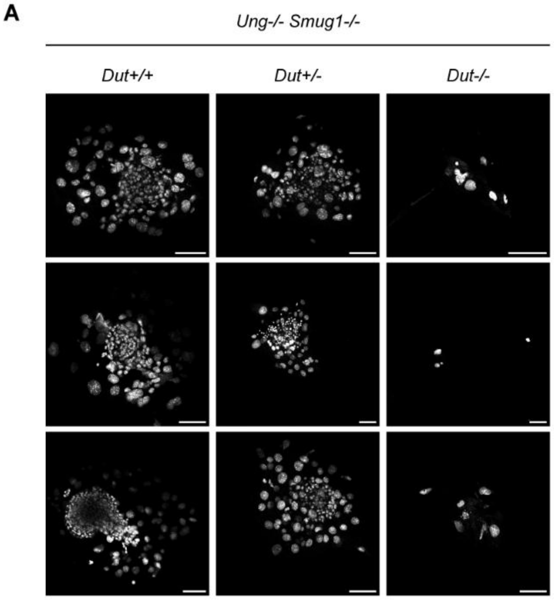


**B**

| Crossing of *Ung-/- Smug1+/+ Dut+/-* mice | | | | | |
|---|---|---|---|---|---|
| Developmental stage | *Dut* genotype | | | Resorbed | No. total |
| | +/+ | +/- | -/- | | |
| 9.5 dpc | 4 | 5 | 0 | NA | 9 |
| Postnatal | 9 | 26 | 0 | NA | 35 |

| Crossing of *Ung+/+ Smug1-/- Dut+/-* mice | | | | | |
|---|---|---|---|---|---|
| Developmental stage | *Dut* genotype | | | Resorbed | No. total |
| | +/+ | +/- | -/- | | |
| 9.5 dpc | 0 | 3 | 0 | NA | 3 |
| Postnatal | 11 | 22 | 0 | NA | 33 |

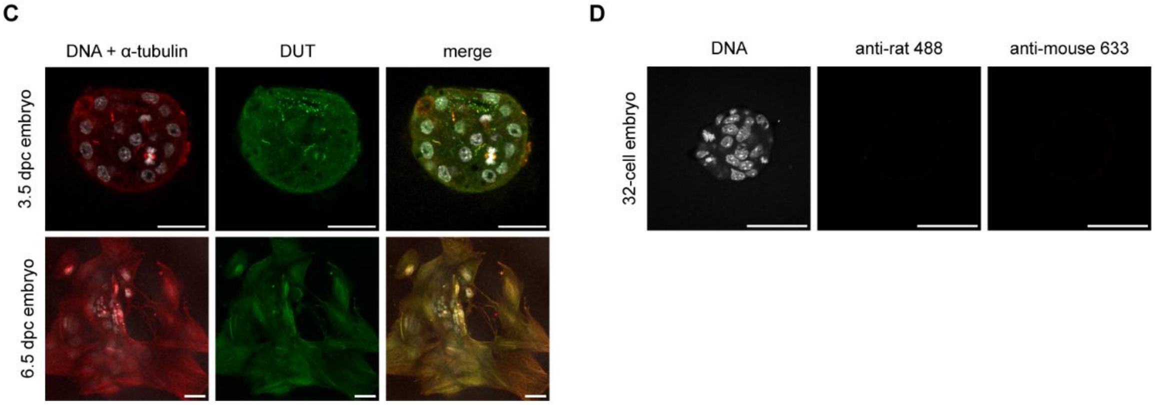


**Supplementary Figure S2. Investigation of *Dut* knockout and WT mouse embryos.** (A) Microscopy analysis of embryos from *Ung*$^{-/-}$ *Smug1*$^{-/-}$ *Dut*$^{+/-}$ crossings at 3.5 dpc. DNA staining is shown in gray. Scale bar 100 µm. (B) Genotyping results of offspring from self-crosses of *Ung*$^{-/-}$ *Smug1*$^{+/+}$ *Dut*$^{+/-}$ or *Ung*$^{+/+}$ *Smug1*$^{-/-}$ *Dut*$^{+/-}$ mice. (C) Immunostaining of DNA (gray), $\alpha$-tubulin (red), and dUTPase (green) in WT 3.5 dpc and 6.5 dpc embryos. Scale bar: 50 µm. (D) Secondary antibody controls using only anti-rat 488 and anti-mouse 633 antibodies. DNA is shown in gray. Scale bar 50 µm.

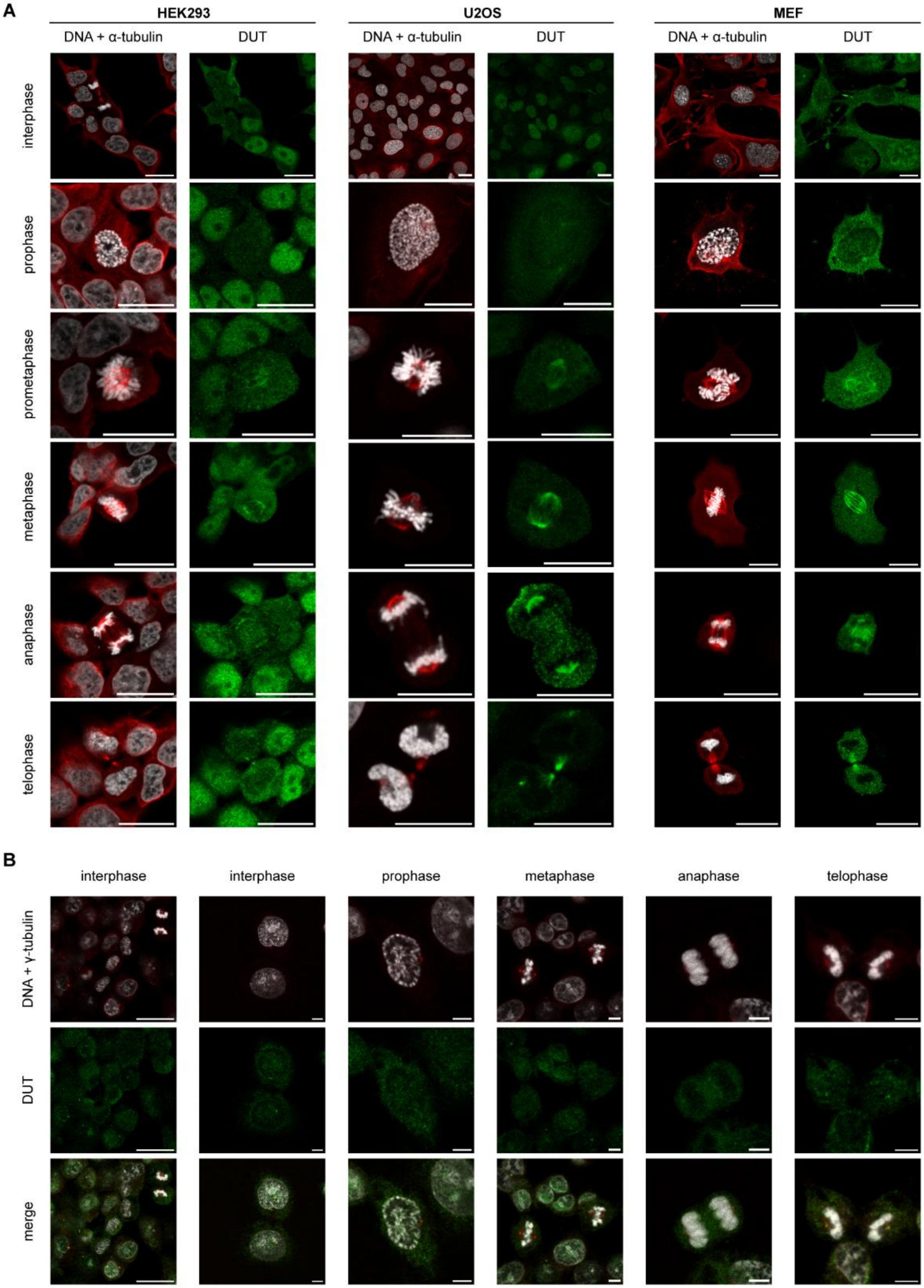
A
HEK293
U2OS
MEF
DNA + α-tubulin
DUT
interphase
prophase
prometaphase
metaphase
anaphase
telophase
B
interphase
prophase
metaphase
anaphase
telophase
DNA + γ-tubulin
DUT
merge

**Supplementary Figure S3. Localization of dUTPase in the different stages of mitosis.** (A) Immunostaining of DNA (gray), α-tubulin (red), and dUTPase (green) in interphase and the different stages of mitosis (prophase, prometaphase, metaphase, anaphase, and telophase) in HEK293, U2OS, and MEF cells. Scale bar 20 µm. (B) Immunostaining of DNA (gray), γ-tubulin (red), and dUTPase (green) in interphase and the different stages of mitosis (prophase, metaphase, anaphase, and telophase) in HCT116 cells. Scale bar on the smaller magnification image is 20 µm, while 5 µm on the close-up images.

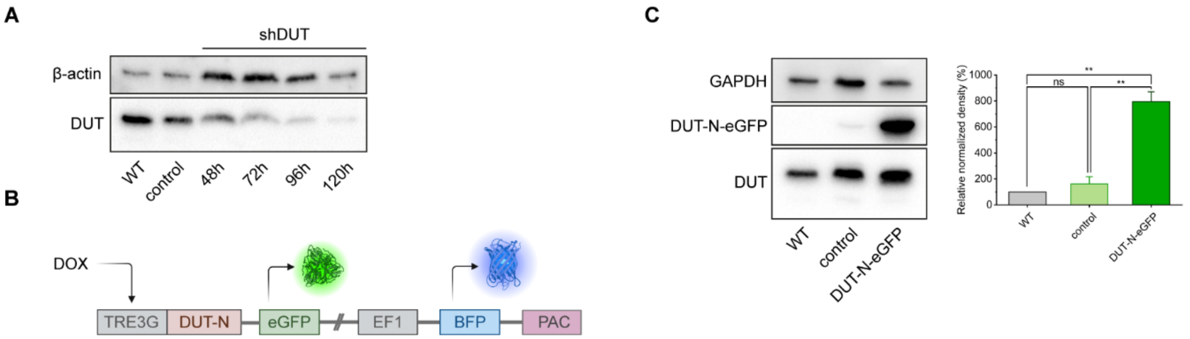


**Supplementary Figure S4. Knockdown and overexpression of dUTPase in human cells.** (A) Western blot analysis of WT, control, and induced shDUT HCT116 cells after 48, 72, 96, and 120 hours. (B) Overexpression cassette contains doxycyclin-inducible (DOX) promoter (TRE3G), nuclear isoform of dUTPase (DUT-N) with eGFP-tag, stably expressing blue fluorescent protein (BFP), and puromycin resistance (PAC) driven by EF1 promoter. (C) Western blot analysis of WT (gray), control (light green), and DUT-N-eGFP overexpressed (green) HCT116 cells. Whiskers indicate standard deviation. Statistical analysis was carried out using One-way ANOVA with Tukey multiple comparisons test (**: $p<0.008$, ns: non-significant).
